# Anks3 mediates cilia dependent polycystin signaling and is essential for adult kidney homeostasis

**DOI:** 10.1101/2025.04.22.649832

**Authors:** Zemeng Wei, Michael Rehman, Jianlei Gu, Gabriel Lerner, Ke Dong, Kasturi Roy, Adrian Cordido, Yiqiang Cai, Xin Tian, Ming Shen Tham, Jean Kanyo, TuKiet T. Lam, Hongyu Zhao, Stefan Somlo

## Abstract

The existence of a cilia-dependent cyst activation (CDCA) pathway underlying autosomal dominant polycystic kidney disease was identified by showing that cyst progression following loss of polycystins is significantly suppressed by removal of structurally intact cilia. We applied translating ribosome affinity purification RNASeq on pre-cystic mouse kidneys to determine a cell-autonomous in vivo translatome associated with CDCA and identified Glis2 as an early effector of polycystin signaling. Here, to discover additional components of CDCA, we used polycystin-dependent Glis2 expression as a functional readout and identified Anks3, from the CDCA pattern translatome, as a candidate cytosolic regulator of polycystin signaling. Anks3 regulates polycystin-dependent Glis2 expression both in vitro and in vivo. Anks3 also undergoes polycystin-dependent changes in phosphorylation state. Inactivation of *Anks3* in *Pkd1* mouse models suppresses cyst progression, but also results in rapidly progressive kidney injury independent of polycystins. *Anks3* inactivation also normalizes a broader spectrum of polycystin-dependent CDCA translatome changes. These findings define Anks3 as a central regulator of cilia dependent polycystin signaling, functioning downstream of cilia and polycystins and upstream of Glis2, and show that Anks3 has broader functions in maintaining renal structural and functional homeostasis.

## Introduction

Autosomal dominant polycystic kidney disease (ADPKD) is the most common monogenic cause of kidney failure. It is most often caused by mutations in either the *PKD1* or *PKD2*, with identified mutations in 78% and 15% of ADPKD families, respectively (1). *PKD1* encodes polycystin-1 (PC1), a complex polytopic integral membrane protein that forms a putative receptor-channel complex in primary cilia with polycystin-2 (PC2), a nonselective cation channel belonging to the transient receptor potential channel family that is encoded by *PKD2* (2, 3). ADPKD presents with cyst formation originating from the epithelia of kidney tubules, with extrarenal manifestations that include bile duct cysts resulting in polycystic liver disease and with an increased risk of intracranial aneurysms that show familial clustering (4-6). While ADPKD is inherited as a dominant trait and is fully penetrant, cyst formation in ADPKD patients is focal, and there are significant intra- and inter-familial phenotypic variability among individuals (7). Cyst initiation results from somatic “second hit” mutations (8-10), the occurrence of which may account for some of the observed variation (11). Other factors including the nature of *PKD* mutations (12), genetic modifiers affecting biogenesis of PC1 (13), and background genetic risk for chronic kidney disease (14) further influence disease severity.

Cysts grow by changing the functional state of kidney tubule epithelial cells including changes in cell shape, transport properties, metabolism and energetics while undergoing low level proliferation (15). These changes promote secondary damaging effects including increased inflammatory responses and tubulointerstitial fibrosis (16). Several signaling pathways and cellular processes have been implicated to be defective in ADPKD, including calcium signaling, cyclic AMP-dependent signaling (17-19), canonical WNT singling (20, 21), mTOR signaling (22, 23), G-protein coupled receptor signaling (24), planar cell polarity (21, 25), cell cycle regulation (26, 27), extracellular matrix organization (28, 29), and cellular metabolic changes (30-35). While there remains a lack of consensus on the precise cell autonomous functions of polycystins, there is general agreement on the importance of primary cilia in the pathogenesis of ADPKD (36-38). The genetic relationship between polycystins and primary cilia has been defined in mouse models of ADPKD. Cyst formation after inactivation of polycystin genes is markedly attenuated by removal of structurally intact cilia (39). We have called the cyst promoting activity of intact cilia devoid of functional polycystins ‘cilia dependent cyst activation’ (CDCA) (37, 40). To begin to understand tubule cell responses to CDCA, we applied translating ribosome affinity purification RNA sequencing (TRAP RNASeq) to pre-cystic mouse kidneys and determined a cell-autonomous translatome specifically associated with CDCA (41). This CDCA pattern translatome is comprised of differentially expressed genes in *Pkd1* knockout with the same direction of change when compared to both wild type and *Pkd1* and cilia double knockouts. These studies led to the discovery that the Krüppel family zinc finger transcription factor Glis2, the causative gene for nephronophthisis type 7 (NPHP7), is an in vitro maker of CDCA activity, an in vivo effector of polycystin and CDCA dependent cyst formation and a therapeutic target in preclinical models of ADPKD (41).

For the current study, we hypothesized that genes whose inactivation suppress *Pkd*-dependent Glis2 upregulation are candidate functional components of the CDCA pathway. Among genes we identified in the CDCA pattern translatome (41), *Anks3* emerged as a compelling candidate. We found that inactivation of *Anks3* suppressed Glis2 upregulation that otherwise results from *Pkd1* inactivation. Anks3, ankyrin-repeat (ANK) and sterile alpha motif (SAM) 3, interacts directly with several ciliopathy proteins including Anks6 (Nphp16) and Nek8 (Nphp9) (42) and indirectly through Anks6 with the Nphp1-4-8 and Nphp2-3-9 modules (43-45). The apparent function of Anks3 as a scaffolding protein for integrating cilia-related signaling and its role in suppressing Glis2 upregulation prompted us to evaluate Anks3 as a candidate regulator of polycystin and cilia related CDCA signaling. We found that *Anks3* functions in CDCA downstream of cilia and polycystins but upstream of Glis2. Polycystins regulate the phosphorylation states of Anks3. *Anks3* knockout in *Pkd1* mouse models suppresses cyst progression but Anks3 has broader functions than CDCA signaling. Anks3 is necessary for maintain adult kidney structural and functional homeostasis and inactivation Anks3 results in diffuse rapidly progressive noncystic kidney injury. The current study defines Anks3 as a central component of the CDCA signaling process in the pathogenesis of ADPKD and as important functional molecule for maintaining renal structural and functional homeostasis.

## Results

### Anks3 regulates Glis2 expression

Our initial report defined a CDCA pattern kidney tubule cell-specific translatome profile determined by TRAP RNASeq in precystic *Pkd1* mouse models induced for gene knockout from postnatal day 28 to 42 (P28-P42) and examined at 7-weeks age (41). We have extended these datasets to include *Pkd1* mouse kidneys at 10-weeks age, and *Pkd2* models at 7- and 10-weeks age. The data are publicly available at the Metabolism and Genomics in Cystic Kidney (MAGICK) web portal (https://pkdgenesandmetabolism.org/). In evaluating ciliopathy related genes other than Glis2 in our TRAP RNASeq datasets, we found that *Anks3* follows the CDCA pattern across all four comparison groups (Figure 1a). In renal primary cells cultured from *Pkd1^fl/fl^; Pax8^rtTA^; TetO^Cre^* mice, cells treated with doxycycline *in vitro* to induce *Pkd1* inactivation (Pkd1^KO^) had 2.5-fold increased *Anks3* mRNA levels compared to “wild type” (WT) cells not treated with doxycycline (Figure 1b). We hypothesized that inactivation of true candidate components of CDCA signaling that function upstream of Glis2 will suppress Glis2 upregulation despite Pkd1^KO^ in a manner analogous to the effects that we had previously shown for cilia inactivation (41). We generated an immortalized cell line using SV40 lentivirus transduction of the Pkd1^KO^ renal primary cells which have high basal levels of Glis2 expression due to *Pkd1* inactivation. We then introduced acute CRISPR inactivation of *Anks3* by nucleofection in these cells. Compared to non-targeting control sgRNA (sgNT), inactivation of Anks3 (sgAnks3) significantly reduced Glis2 mRNA and protein expression (Figure 1c-e; Supplementary Figure 1). The efficiency of *Anks3* knockout was confirmed by Sanger sequencing and by immunoblotting using a polyclonal antibody against Anks3 that we developed (Figure 1d; Supplementary Figure 1). To improve in vivo and in vitro assessment of Glis2 responses, we made a Glis2-HaloTag7 (*Glis2^HT^*) knockin mouse in which HaloTag7 is inserted in-frame before the termination codon of *Glis2* (Supplementary Figure 2). Homozygous *Glis2^HT/HT^* mice (noted as Glis2^Halo^) are normal without a nephronophthisis phenotype indicating the fusion protein is fully functional (Supplementary Figure 2). Kidney primary cell cultures from *Glis2^HT/HT^; Pkd1^fl/fl^; Pax8^rtTA^; TetO^Cre^* mice were treated with doxycycline in vitro and immortalized with SV40 lentivirus to produce Glis2^Halo^+Pkd1^KO^ cell lines. Live cells labeled with tetramethylrhodamine (TMR) HaloTag ligand show nuclear expression of Glis2^Halo^ by immunofluorescence in the presence of sgNT (Figure 1f). Acute CRISPR knockout of *Anks3* (sgAnks3) results markedly reduced Glis2^Halo^ nuclear expression (Figure 1f). We isolated single-cell clones to obtain a pure population of complete *Anks3* knockout cells (Glis2^Halo^+Pkd1^KO^+Anks3^KO^). Stable transfection of triple-HA epitope tagged Anks3-HA showed that re-expressing Anks3 in a *Pkd1* and *Anks3* null background upregulated Glis2 nuclear expression (Figure 1g,h). These data show that *Anks3* regulates Glis2 expression in *Pkd1* knockout condition in vitro and support a hypothesized causal relationship between Anks3 and Glis2 upregulation in the polycystin-dependent CDCA pathway.

**Figure 1.**
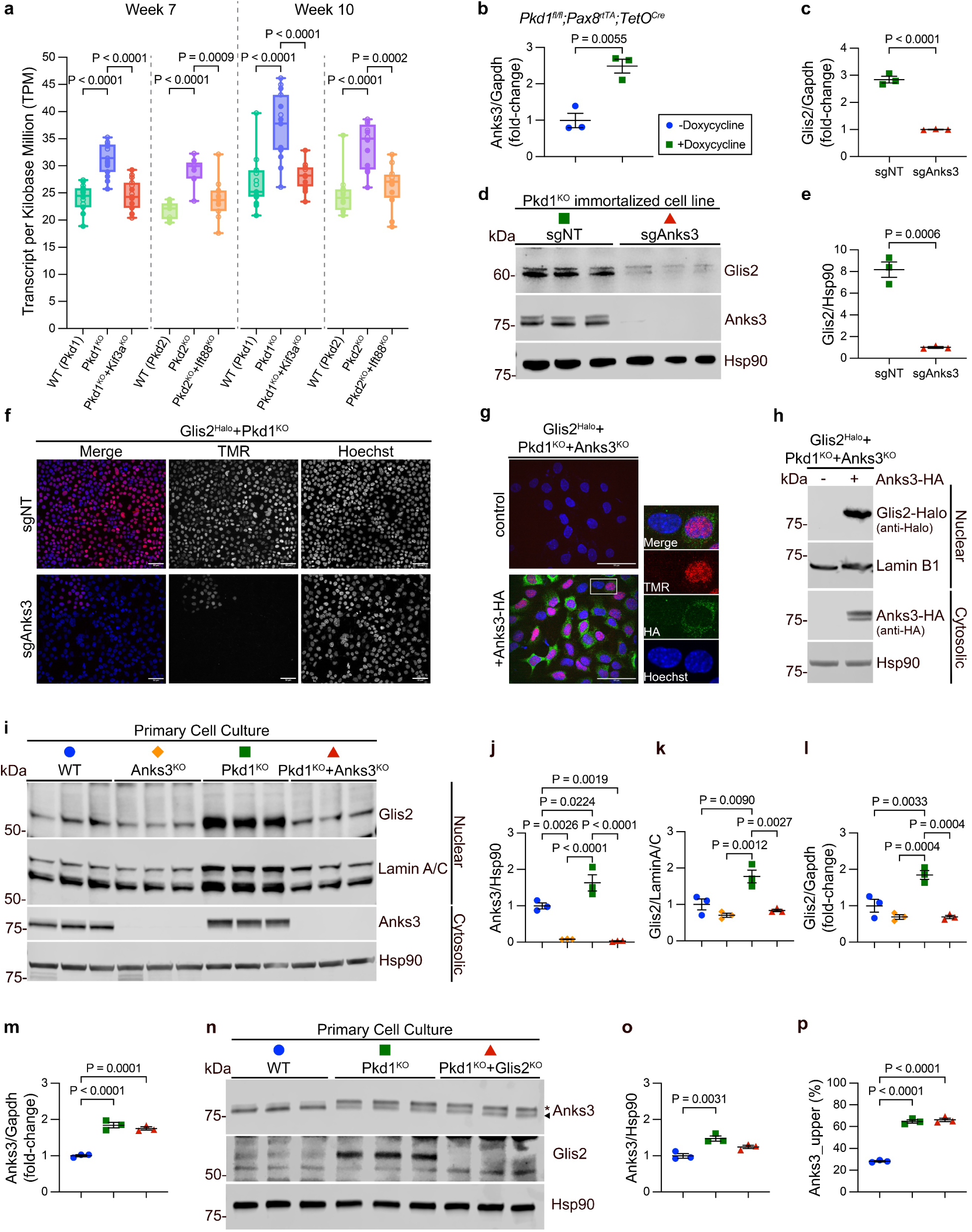
Anks3 regulates polycystin-dependent Glis2 expression. **a**, *Anks3* expression from in vivo TRAP RNASeq (https://pkdgenesandmetabolism.org/expression/Anks3) shown as combined normalized TPM values from male (closed symbols) and female (open symbols) samples. Multiple-group comparisons for each genotype-timepoint group were performed by one-way ANOVA with Tukey’s multiple-comparison test. **b**, *Anks3* is upregulated following *Pkd1* inactivation. RT-qPCR of primary culture of kidney cells from *Pkd1^fl/fl^; Pax8^rtTA^; TetO^Cre^* mice with and without in vitro doxycycline induced *Pkd1* inactivation. Data points represent independent primary cell cultures from three mice. **c-e**, Anks3 knockout suppresses Glis2 upregulation in Pkd1^KO^ cells with acute CRISPR gene targeting by non-targeting (sgNT) or Anks3 (sgAnks3) guide RNAs. **c**, RT-qPCR, **d**, immunoblots, and **e**, quantification of whole cell lysates. **c,e**, Fold changes for **c**, *Glis2* mRNA normalized to *Gapdh* and **e**, Glis2-to-Hsp90 ratio for each lane in **d** are shown relative to the mean in sgAnks3 group, which is set to 1.0. Statistical significance by unpaired two-tailed Student’s *t* test presented as mean ± s.e.m. **f**, Nuclear Glis2^Halo^ expression in Glis2^Halo^+Pkd1^KO^ cell lines with (sgAnks3) and without (sgNT) acute CRISPR knockout of *Anks3* live labelled with TMR, followed by fixation and staining with Hoechst. **g,h**, Re-expression of Anks3-HA upregulates nuclear Glis2^Halo^. **g**, Glis2^Halo^+Pkd1^KO^+Anks3^KO^ cells with stable transfection of Anks3-HA or untransfected controls live labeled with TMR, followed by fixation and anti-HA staining. **h**, Immunoblots of nuclear and cytosolic fractions of cells from **g**. **f,g**, Scale bar, 50 μm. **i-l**, Glis2 suppression in an allelic series of primary kidney cell cultures from *Pkd1* and *Anks3* mutant mice. **i**, Immunoblots and **j,k**, quantifications of cytosolic (**j**) and nuclear (**k**) fractions. Cells were treated with doxycycline in vitro to induce gene deletions. Fold changes in **j** are the ratio of the sum of two Anks3 bands to Hsp90 and in **k** the ratio of Glis2 to Lamin A/C; ratios are shown relative to the means in WT group, which is set to 1.0. **l**, RT-qPCR of primary culture of kidney cells from **i** with fold changes for *Glis2* normalized to *Gapdh* and are shown relative to the mean in WT group, which is set to 1.0. **m-p**, Pkd1 dependent changes in Anks3 are not regulated by Glis2. **m**, RT-qPCR, **n**, immunoblots and **o,p**, quantifications from primary kidney cell culture lysates with the indicated genotypes. All cells were treated with doxycycline in vitro to induce respective gene deletions. **m**, Fold changes for *Anks3* mRNA expression normalized to *Gapdh* and shown relative to the mean in WT group, which is set to 1.0. **o**, Fold changes for ratio of the sum of two Anks3 bands to Hsp90, shown relative to the mean of the ratio in WT group, which is set to 1.0. **p**, The ratio of densitometric volume of the upper band of Anks3 to sum of both bands of Anks3 expressed as a percentage. **j-m,o,p**, Multiple-group comparisons were performed by one-way ANOVA followed by Tukey’s multiple-comparison test, presented as mean ± s.e.m.

We produced a conditional *Anks3^fl^* allele (Supplementary Note; Supplementary Figure 3) and generated mice with the following four genotypes: wild type (WT), *Anks3^fl/fl^; Pax8^rtTA^; TetO^Cre^* (Anks3^KO^), *Pkd1^fl/fl^; Pax8^rtTA^; TetO^Cre^* (Pkd1^KO^), and *Anks3^fl/fl^; Pkd1^fl/fl^; Pax8^rtTA^; TetO^Cre^* (Pkd1^KO^+Anks3^KO^) mice. Primary cell cultures from kidneys of mice with each genotype were treated with doxycycline to induce the respective gene knockouts in vitro and nuclear and cytosolic fractions were prepared. Anks3 and Glis2 protein expression in the cytosolic and nuclear fractions, respectively, was increased in Pkd1^KO^ compared to WT (Figure 1i-k). Anks3 appears as a doublet and the sum of the doublet bands was used in quantitation (Figure 1i,j). Of note, there is an increase in the Anks3 upper band relative to the lower band in Pkd1^KO^ compared to WT (Figure 1i). This is further investigated in studies described below. There was complete knockout of Anks3 in the cytosolic fractions (Figure 1i,j) and nuclear Glis2 was significantly reduced in Pkd1^KO^+Anks3^KO^ double inactivation compared to Pkd1^KO^ (Figure 1i,k). RT-qPCR showed that the suppression of Glis2 by Anks3 inactivation occurs at the level steady state mRNA expression (Figure 1l). Of note, the Anks3^KO^ showed a non-significant trend toward Glis2 transcript and protein levels below that seen in WT (Figure 1i,k,l) raising the possibility that Anks3 is a regulator of Glis2 expression even without *Pkd1* inactivation. Finally, since Glis2 is a transcription factor upregulated by CDCA, we determined whether Anks3 is a transcriptional target of Glis2 upregulation. We made primary kidney cell cultures form WT, *Pkd1^fl/fl^; Pax8^rtTA^; TetO^Cre^*, and *Glis2^fl/fl^; Pkd1^fl/fl^; Pax8^rtTA^; TetO^Cre^* mice and treated them in vitro with doxycycline to produce WT, Pkd1^KO^ and Pkd1^KO^+Glis2^KO^ cells, respectively (Figure 1m-p). Pkd1^KO^ showed upregulation of Anks3 mRNA expression (Figure 1m) and total protein (Figure 1n,o). There was also a shift to more of the upper band for Anks3 (Figure 1p). These changes in Anks3 expression persisted in Pkd1^KO^+Glis2^KO^ cells indicating that the polycystin dependent changes in Anks3 expression are not regulated by Glis2. In aggregate, the in vitro data show that Anks3 functions downstream of polycystins and upstream of Glis2 to regulate Glis2 transcript and protein expression in a polycystin and CDCA dependent manner.

### Polycystins regulate the phosphorylation state of Anks3

Immunoblotting of Anks3 showed two bands with intensities that vary with Pkd genotype (Figure 1i,n,p). The two bands were also detected using anti-V5 and anti-HA antibodies with overexpressed Anks3 cDNA with N-terminal V5 and C-terminal HA tags (Supplementary Figure 4a). This excluded alternative splicing and protein cleavage as the source of the two bands. There is evidence that Anks3 is phosphorylated (https://www.phosphosite.org/homeAction). When lysates from IMCD3 cells expressing V5-Anks3-HA were treated with lambda protein phosphatase, the upper band of Anks3 completely disappeared indicating that differential phosphorylation states account for the multiple bands of Anks3 seen on immunoblots (Supplementary Figure 4b). We next cultured kidney cells from *Pkd1^fl/fl^; Pax8^rtTA^; TetO^Cre^* mice and confirmed increased prominence of the upper band of endogenous Anks3 following Pkd1^KO^ after doxycycline treatment (Figure 2a). The upper bands in both WT and Pkd1^KO^ disappeared after treatment with lambda protein phosphatase (Figure 2a). To determine whether the polycystin dependent changes in Anks3 expression level and phosphorylation state occurred in vivo, we applied the Anks3 antibody to whole kidney tissue lysates of 19-week-old WT and cystic Pkd1^KO^ mice (Figure 2b). We detected both an increase of total Anks3 protein (Figure 2c), and a significantly elevated level of the upper phosphorylated Anks3 band (Figure 2d). There was a similar shift to the upper phosphorylated Anks3 band in whole kidney lysates of *Pkd2^fl/fl^; Pax8^rtTA^; TetO^Cre^* (Pkd2^KO^) mice (Figure 2e-g). We next determined whether re-expression of *Pkd1* or *Pkd2* restored phosphorylation state of Anks3 to the WT pattern. To do this, we cultured primary kidney cells from our *Pkd1* and *Pkd2* re-expression mouse models (46), *Pkd1^fl/FSF^; Pax8^rtTA^; TetO^Cre^; Rosa^FlopER^* mice and *Pkd2^fl/-^; Pkd2^FSF^; Pax8^rtTA^; TetO^Cre^; Rosa^FlopER^*, respectively. The cells either received no treatment (WT), doxycycline treatment alone (Pkd1^KO^ or Pkd2^KO^), or doxycycline treatment followed by 4-OH-tamoxifen treatment (Pkd1^re-expression^ or Pkd2^re-expression^; Figure 2h). Immunoblot using whole cell lysates showed a significant increase in total Anks3 protein levels and a relative increase of upper phosphorylated species in the Pkd1^KO^ and Pkd2^KO^ conditions (Figure 2i-n). Re-expression of *Pkd1* and *Pkd2* in the respective knockouts restored the phosphorylation state of Anks3 to the WT pattern (Figure 2i-n). We performed Phos-tag electrophoresis to better separate the polycystin-dependent phosphorylated states of Anks3 in the *Pkd1* re-expression cell models. Phos-tag resolved at least three bands, with the lower two present in WT and a shift of most of Anks3 to a novel upper third band in the Pkd1^KO^ (Figure 2o). There was reversion to the lower two bands with a minor persistence of the upper third band following re-expression of *Pkd1*. In addition, we confirmed the polycystin dependent modulation of Anks3 phosphorylation state in vivo in whole kidney tissue lysates from *Pkd1* and *Pkd2* re-expression mouse models (Figure 2p,q). Overall, the data show that Anks3 undergoes multiple phosphorylation steps and that polycystin-1 and polycystin-2 regulate the phosphorylation states of Anks3.

**Figure 2.**
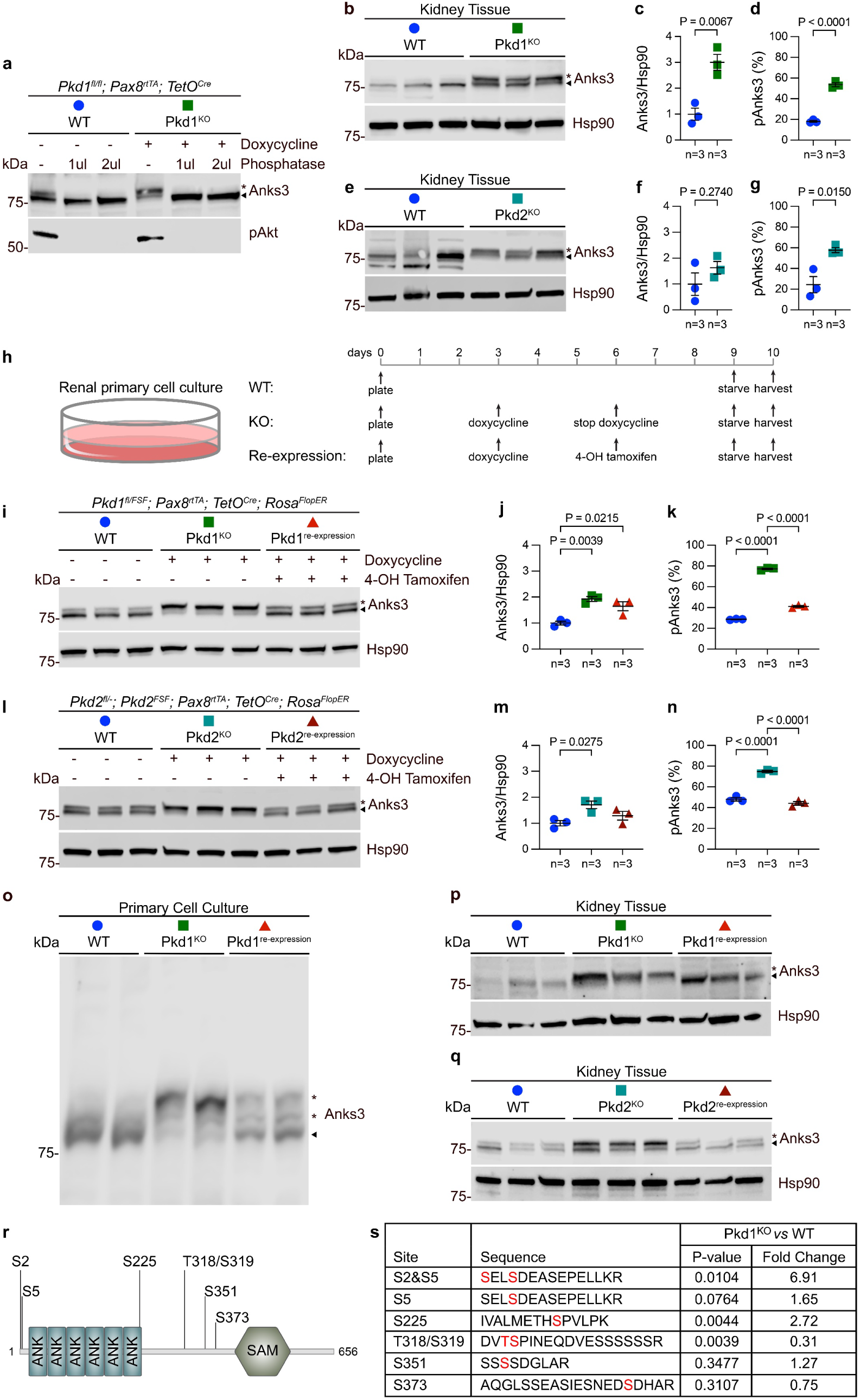
Polycystins regulate the phosphorylation state of Anks3. **a**, Immunoblots of Lambda protein phosphatase treated kidney primary cell lysates with and without in vitro doxycycline induction. Phospho-Akt (pAkt) serves as positive control for phosphatase activity. **b-g**, Immunoblots (**b,e**) and quantifications (**c,d,f,g**) of kidney tissue lysates from 19 week old Pkd1^KO^ (**b-d**) and Pkd2^KO^ (**e-g**) mice. **h-o**, Anks3 phosphorylation state is reversible in a polycystin dependent manner. **h**, Schematic for the in vitro inactivation (doxycycline) and re-expression (4-OH tamoxifen) of polycystin in primary kidney cell cultures. **i-n**, Immunoblots (**i,l**) and quantifications (**j,k,m,n**) of Anks3 expression and phosphorylation state in *Pkd1* (**i-k**) and *Pkd2* (**l-n**) models. **o,** Lysates of first two lanes of each genotype group in **i** were run on Phos-tag gel further resolve phosphorylated forms. **c,f,j,m**, Fold changes of densitometric ratio of the sum of two Anks3 bands to Hsp90 relative to the mean of the ratio in WT group, which is set to 1.0. **d,g,k,n**, The ratio of densitometric volume of the upper phosphorylated band of Anks3 to total of both bands expressed as a percentage. Three mice per genotype were used in **b,e,p,q** and three independent cell cultures from a single mouse of each genotype were used in **i,l**. **c,d,f,g**, Statistical significance was determined by unpaired two-tailed Student’s *t* test and presented as mean ± s.e.m. **j,k,m,n**, Multiple-group comparisons were performed by one-way ANOVA followed by Tukey’s multiple-comparison test, presented as mean ± s.e.m. **p,q**, Immunoblots of total kidney lysates from 19 week *Pkd1* (**p**) and *Pkd2* (**q**) mouse models probed with anti-Anks3 after doxycycline induced knockout (Pkd1^KO^ and Pkd2^KO^) and following tamoxifen induced re-expression (Pkd1^re-expression^ and Pkd2^re-expression^). Hsp90 was used as loading control. **r**, Schematic of Anks3 protein with labeled functional domains (ankyrin repeats, ANK; sterile alpha motif, SAM) and phosphorylation sites. **s**, Quantitative analysis of normalized abundance of specific phosphorylated Anks3 peptides. Statistical significance was determined by unpaired two-tailed Student’s *t* test on log_2_ transformed normalized abundance.

We next sought to determine the specific sites in Anks3 that result in the differential phosphorylation in the Pkd1^KO^ condition. We performed liquid chromatography-tandem mass spectrometry (LC-MS/MS) on immunoprecipitates of Anks3 from triplicate samples of renal primary cell cultures from *Pkd1^fl/fl^; Pax8^rtTA^; TetO^Cre^* mice, treated with (Pkd1^KO^) or without (WT) doxycycline. We first confirmed that all forms of Anks3 can be immunoprecipitated and enriched by the Anks3 antibody (Supplementary Figure 4c). Mass spectrometric analyses showed 74% and 61% coverage of Anks3 in WT and Pkd1^KO^ samples, respectively (Supplementary Figure 5a,b). We identified six phosphorylation sites of Anks3 present in both samples, including Ser2, Ser5, Ser225, Thr318/Ser319, Ser351, and Ser373. The location of phosphorylation sites relative to the functional domains of Anks3 is shown in Figure 2r. There was no novel phosphorylation site detected specifically in Pkd1^KO^ samples. There was, however, a 6.9-fold increase in Ser2 and Ser5 dual-phosphorylated peptides in Pkd1^KO^ compared to WT, while peptides with phosphorylation at Ser5 alone were not significantly different based on genotype (Figure 2s). The MS/MS spectrum of single- and dual-phosphorylated Anks3 peptide containing Ser2 and Ser5 are shown in Supplementary Figure 5c. Notably, these are the N-terminal peptides of the full length Anks3 protein. The first methionine is removed from the N-terminus and Ser2 is acetylated in all forms of Anks3 regardless of the phosphorylation state or *Pkd1* genotype. In addition, phosphorylation at Ser225 was significantly increased by 2.7-fold in Pkd1^KO^ compared to WT and phosphorylation at Thr318/Ser319 was significantly decreased in Pkd1^KO^ (Figure 2s). Phosphorylation at Ser351 and Ser373 were not change significantly in Pkd1^KO^ compared to WT. The findings are consistent with the immunoblot and Phos-tag data showing multiple phosphorylation states with changes in the relative intensity of these phosphorylation states based on Pkd genotype. Polycystin function regulates phosphorylation of Anks3 primarily at Ser2 but also has significant effects at Ser225 and Thr318/Ser219.

### Anks3 inactivation suppresses polycystic kidney disease but results in diffuse tubular injury

We next applied the four allele combinations WT, Anks3^KO^, Pkd1^KO^, and Pkd1^KO^+Anks3^KO^ to evaluate the effect of *Anks3* inactivation on progression of polycystic kidney disease in vivo. We first used an early-onset model of ADPKD in which doxycycline water was administrated to the nursing dams from postnatal day 0 (P0) to postnatal day 14 (P14) and kidneys from the pups were examined at P14. Compared to the Pkd1^KO^ which developed substantial polycystic kidney disease at P14, the Pkd1^KO^+Anks3^KO^ double mutants were significantly protected from cyst progression as indicated by normal kidney weight to body weight ratio, cystic index, and blood urea nitrogen (BUN) (Figure 3a-e; Supplementary Figure 6). Age-matched doxycycline-induced Anks3^KO^ mice did not show overt histopathological changes by light microscopy or changes in kidney function (Figure 3b,e). Consistent with the in vitro studies, the significant upregulation of Glis2 expression in the nuclear fraction of Pkd1^KO^ kidney tissue lysates was suppressed to WT levels in Pkd1^KO^+Anks3^KO^ kidneys at P14 (Figure 3f-g). These findings show that inactivation of *Anks3* suppresses cyst growth in early-onset model of ADPKD and this correlates with normalization of nuclear Glis2 expression.

**Figure 3.**
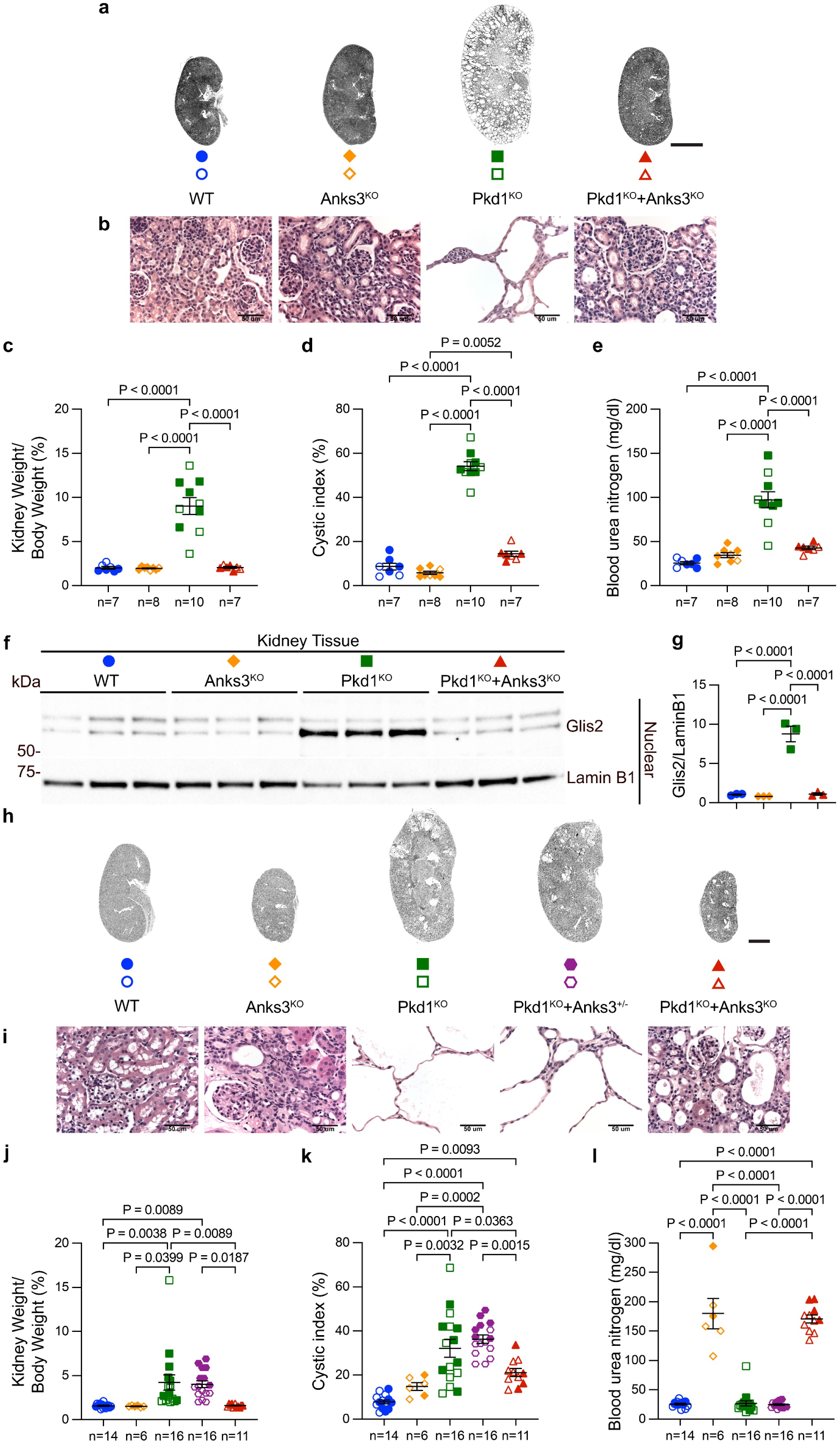
Anks3 inactivation suppresses cyst growth but results in kidney injury. **a-e,** *Anks3* inactivation suppresses cyst formation in an early-onset ADPKD model. **a**, Representative images of kidneys from mice at P14. Oral doxycycline was administrated to the nursing dams from P0-P14, and the kidneys from the pups were examined at P14. **b**, Representative images of H&E staining for the corresponding genotypes. **c-e**, Aggregate quantitative data for kidney-to-body weight ratio (**c**), cystic index (**d**), and blood urea nitrogen (**e**). Colors and symbol shapes correspond to genotype defined in **a**; closed symbols, male; open symbols, female. *n*, number of mice in each group. **f,g**, Anks3 inactivation suppresses Glis2 upregulation in Pkd1^KO^ kidney tissues. **f**, Immunoblots and **g**, quantifications of nuclear fraction of whole kidney tissue from mice in **a** probed with anti-Glis2. Fold changes in ratio of Glis2 to Lamin B1 for each lane are shown relative to the mean in wild type (WT) group, which is set to 1.0. **h-l**, Anks3 inactivation reduces cyst growth but results in kidney damage in an adult-onset model. **h**, Representative images of kidneys from mice at 13 weeks of age. All mice were administrated with oral doxycycline from P28-P42 and were examined at 13 weeks. **i**, Representative images of H&E staining for the corresponding genotypes. **j-l**, Aggregate quantitative data for kidney-to-body weight ratio (**j**), cystic index (**k**), and blood urea nitrogen (**l**). Colors and symbol shapes correspond to genotype defined in **h**; closed symbols, male; open symbols, female. *n*, number of mice in each group. **c-e,g,j-l**, Multiple-group comparisons were performed by one-way ANOVA followed by Tukey’s multiple-comparison test, presented as mean ± s.e.m. **a,h**, Scale bar, 2 mm. **b,i**, Scale bar 50 μm.

We next investigated the effect of *Anks3* inactivation on polycystic kidney disease progression in an adult-onset mouse model of ADPKD (Figure 3h-l). All experimental mice were administrated doxycycline in drinking water from P28 to P42 and were examined at 13-weeks age. Compared to the Pkd1^KO^ cystic controls, Pkd1^KO^+Anks3^KO^ mice were significantly protected from kidney cyst growth (Figure 3h-l; Supplementary Figure 7) when evaluated by kidney weight to body weight ratio (Figure 3j) and cystic index (Figure 3k). However, while cyst progression was suppressed, both Anks3^KO^ single mutant and Pkd1^KO^+Anks3^KO^ double mutant mice had significantly increased BUN (Figure 3l) indicating disruption of renal function. Given the abnormal kidney function associated with Anks3^KO^, we considered whether inactivation of a single copy of *Anks3* has any protective effect on cyst progression. *Anks3^fl/+^; Pkd1^fl/fl^; Pax8^rtTA^; TetO^Cre^* (Pkd1^KO^+Anks3^+/–^) mice do not show improvement of cyst progression at 13 weeks (Figure 3h-l). There was no reduction in polycystic kidney disease progression when we extended observation of Pkd1^KO^+ Anks3^+/–^ mice to 19 weeks (Supplementary Figures 8, 9), indicating that haploinsufficiency of *Anks3* dosage is not sufficient to suppress cyst progression.

We examined the histopathologic kidney phenotype resulting from Anks3^KO^ in further detail. Anks3^KO^ mice with knockout induced from P28-P42 developed progressive kidney failure from 7 to 13 weeks age, with rising BUN associated with decreasing body weight and kidney weight (Supplementary Figure 10a-d). The 13-week endpoint was used for the adult models because both Anks3^KO^ and Pkd1^KO^+Anks3^KO^ mice have reduced survival beyond this timepoint. At 7-weeks, Anks3^KO^ mice still had normal body weight, kidney weight, and BUN compared to WT controls (Supplementary Figure 10a-d). Hematoxylin and eosin (H&E), Masson-trichrome, and periodic acid-Schiff (PAS) staining, showed normal glomerular and tubulointerstitial appearance (Figure 4a,b; Supplementary Figure 10e,f). By 10-weeks age, Anks3^KO^ mice had significantly elevated BUN indicative of impaired renal function (Supplementary Figure 10a-d). Histological examination showed a diffuse tubular injury pattern with luminal dilation and moderate-to-severe interstitial fibrosis, tubular atrophy, and interstitial infiltration of lymphocytes (Figure 4c,d; Supplementary Figure 10g,h). At 13-weeks, body weight, kidney weight and BUN of Anks3^KO^ mice were all significantly altered compared to WT (Supplementary Figure 10a-d), and the histological changes progressed (Figure 4e,f; Supplementary Figure 10i,j). Histopathological features included signs of interstitial inflammation (Figure 4d,g), perivascular lymphocytic aggregates (Figure 4f,h), mitotic figures (Figure 4f,i), tubule cell vacuolization (Figure 4f,j), loss of brush border (Figure 4k; Supplementary Figure 10j), and presence of uromodulin casts (Figure 4l; Supplementary Figure 10j). Immunostaining of the proximal tubule marker megalin on cryosections of 7-, 10-, and 13-week Anks3^KO^ kidneys showed progressive tubule atrophy and dropout compared to WT (Figure 4a-f).

**Figure 4.**
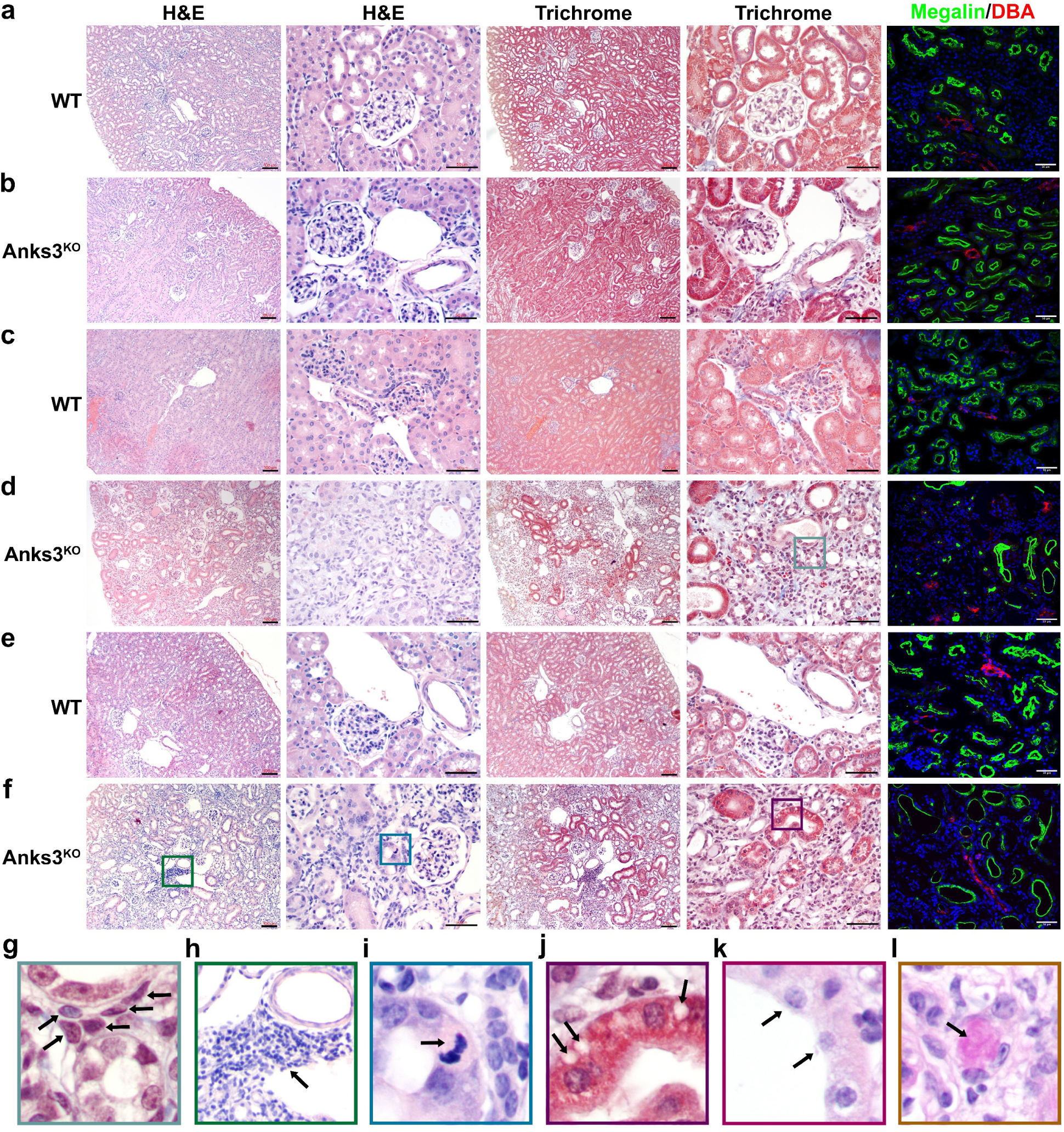
Kidney tubule selective inactivation of Anks3 results in progressive kidney damage. **a-f**, Representative images from indicated genotypes (WT, Anks3^KO^) of hematoxylin and eosin (H&E) and Masson-trichrome (Trichrome) stained kidney histological sections at low (*left*) and high (*right*) magnification for each stain; immunofluorescence images of megalin and dolichos biflorus agglutinin (DBA) to stain proximal tubules and collecting ducts, respectively. **a,b**, 7-weeks; **c,d**, 10-weeks; **e,f**, 13-weeks. Scale bar, 100 μm (low magnification H&E, Trichrome) and 50 μm (high magnification and immunofluorescence). **g-l**, Enlarged areas from panels **d,f** above (**g-j**) and Supplementary Figure 10j (**k,l)** with the corresponding border colors to show specific histological features: interstitial inflammation (**g**; arrows, mononuclear cells); perivascular lymphocytic aggregate (**h**; arrow), mitotic figure (**i**; arrow), tubular cell vacuolization (**j**; arrows), loss of brush boarder (**k**; arrows), and presence of uromodulin casts (**l**; arrow).

We next determined whether Anks3 function in the kidney is dependent on cilia. Anks3 is associated with ciliopathy phenotypes including nephronophthisis and left-right axis formation during development (42, 45, 47, 48). While an initial report suggested Anks3 may be expressed in primary cilia (49), subsequent studies show Anks3 is confined to the cytoplasm (50, 51). We examined the subcellular localization of Anks3 by stable expression of a single plasmid construct of Anks3-EGFP and the Nphp3^1-200^-mApple cilia marker (41) linked by a tandem P2A-T2A peptide in WT and Pkd1^KO^ LLC-PK_1_ cells. Live cell confocal imaging of native fluorescence showed Anks3 localized only in the cytosol and not in primary cilia, both in presence and absence of PC1 (Supplementary Figure 11a). Primary kidney cell cultures from *Anks3^fl/fl^; Pax8^rtTA^; TetO^Cre^* mice with or without doxycycline induced inactivation of Anks3 showed Arl13b and Pkd2 expression in cilia and Anks6 and Invs expression in the inversin compartment of cilia in both WT and Anks3^KO^ conditions (Supplementary Figure 11b-d). We next evaluated whether the kidney injury phenotype of Anks3^KO^ is dependent on structurally intact cilia. *Anks3^fl/fl^; Ift88^fl/fl^; Pax8^rtTA^; TetO^Cre^* (Anks3^KO^+Ift88^KO^) mice and their littermate WT controls were given doxycycline water from P28 to P42 and were examined at 10-weeks age. Anks3^KO^+Ift88^KO^ mice displayed similar diffuse tubular injury phenotype as Anks3^KO^ single mutant and showed significantly elevated BUN compared to wild type (Supplementary Figure 12a-d), showing that inactivation of Ift88 cannot ameliorate the phenotype of Anks3^KO^. Anks3 inactivation results in disruption of kidney structural and functional homeostasis and this effect is not dependent on the presence of intact primary cilia structure.

Tubular injury often results in a regenerative response marked by proliferation of surviving cells. We assessed tubule cell proliferation in 7- and 10-week Anks3^KO^ kidneys by Ki67 staining (Figure 5a-c). As early as 7-weeks, when the kidney histology is largely normal (Figure 4b), inactivation of *Anks3* resulted in a significant increase in proliferation in proximal tubules and a trend toward increased proliferation in collecting ducts (Figure 5b,c). Inflammatory responses and fibrosis are also hallmarks in kidney injury. RT-qPCR from 7-week Anks3^KO^ kidneys showed significant increases in F4/80 macrophage marker mRNA levels in addition to significantly elevated mRNA levels for T cell markers (CD3G, CD3E) comprised of both helper CD4+ T cells (CD4) and cytotoxic CD8+ T cells (CD8A), as well as natural killer NK cells (CD161) (Figure 5d). B cells (CD20) were not significantly increased. RT-qPCR at 7 weeks also showed activation of myofibroblasts with significantly increased platelet-derived growth factor receptor β (PDGFRβ) mRNA (Figure 5d). These mRNA expression patterns were correlated with immunofluorescence analyses showing F4/80 macrophage infiltration initially in the cortex at 7-weeks but extending to the medulla by 10 weeks (Figure 5e). A similar pattern of progressive interstitial inflammation initially was seen with the cytokine tumor necrosis factor-α (TNFα) (Figure 5f). Immunostaining of PDGFRβ and α-smooth muscle actin (α-SMA) suggested the early occurrence and subsequent progression of tubulointerstitial myofibroblast activation in Anks3^KO^ kidney injury (Figure 5g-h). Loss of Anks3 in renal epithelial cells triggers progressive tubule injury associated with tubulointerstitial inflammation and fibrosis. In addition to its CDCA activity regulating Glis2 expression downstream of polycystin signaling, Anks3 is an essential component maintaining normal kidney tubule epithelium homeostasis.

**Figure 5.**
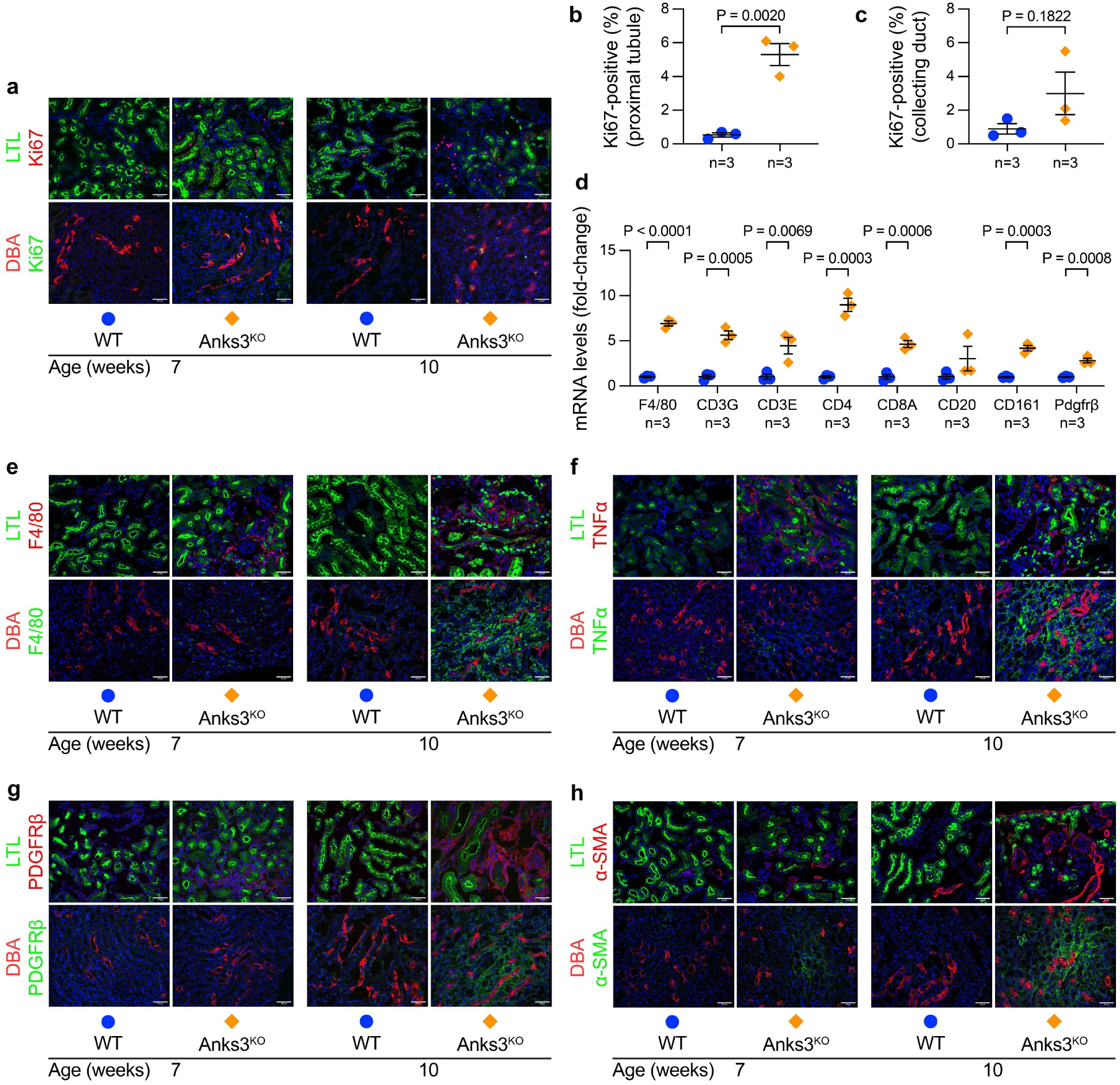
Progressive kidney damage following adult inactivation of *Anks3*. **a,e,f,g,h**, Immunofluorescence of Ki67 (**a**), F4/80 (**e**), TNFα (**f**), PDGFRβ (**g**) and αSMA (**h**) on cryosections of the indicated genotypes and ages. Lotus tetragonolobus lectin (LTL) and DBA were used as tubule segment markers for proximal tubule and collecting duct, respectively. Upper panels, cortex; lower panels, medulla. Scale bar, 50 μm. **b,c**, Quantifications of Ki67-postive nuclei in at least 1000 proximal tubule (**b**) and collecting duct (**c**) nuclei per sample. **d**, RT-qPCR of whole kidneys from mice at 7 weeks age. Fold changes for F4/80, CD3γ, CD3ε, CD4, CD8α, CD20, CD161 and Pdgfrβ mRNA expression are normalized to *Gapdh* and are shown relative to the mean in WT group, which is set to 1.0 for each gene. **b,c,d**, Statistical significance was determined by unpaired two-tailed Student’s *t* test and presented as mean ± s.e.m. *n*, number of mice in each group.

### Anks3 has separate roles in CDCA signaling and kidney homeostasis

Finally, we determined the relationship between the transcriptional responses following *Anks3* and *Pkd1* inactivation. Specifically, we sought to address two questions: *i*) what are the global transcriptomic changes resulting from inactivation of *Anks3*; and *ii*) what are the responses of the other CDCA pattern genes following loss of *Anks3* in the setting of *Pkd1* inactivation. To do this, we performed RNASeq across four genotypes, WT, Anks3^KO^, Pkd1^KO^, and Pkd1^KO^+Anks3^KO^, in two biological systems, primary kidney cell cultures and bulk kidney tissues. All samples were from male mice to limit confounding by sex-dependent gene expression changes. The use of both cell culture and kidney tissue systems was expected to provide some resolution between cell-autonomous and whole organ responses. The primary kidney cell cultures were treated with doxycycline in vitro to induce the respective gene inactivation and grown to form cilia prior to processing for RNASeq. For kidney tissue RNASeq, mice were induced with doxycycline from P28-P42 and RNA was prepared at 7 weeks age. This timepoint antecedes overt histological evidence of kidney injury or cyst formation in the Anks3^KO^ (Figures 4, 5) and Pkd1^KO^ models, respectively, and corresponds to the timepoint used in the TRAP RNASeq studies (41). All samples underwent library construction and sequencing together to limit batch effects. Principal component analysis (PCA) showed adequate clustering of samples by genotype in both cell and tissue samples (Supplementary Figure 13a,b). In primary cell cultures, Pkd1^KO^ clustered separately from the other genotypes, whereas in kidney tissue, the two genotypes that contained Anks3^KO^ clustered separately from WT and Pkd1^KO^ (Supplementary Figure 13a,b). These patterns suggest that Pkd1^KO^ shows greater cell autonomous change in cultured cells after knockout, whereas Anks3^KO^ induces a significant non-cell autonomous response in whole kidney at a stage when Pkd1^KO^ responses are still relatively modest.

We explored this hypothesis further by examining differentially expressed genes (DEG) in five pairwise comparisons across the four genotypes (Supplementary Figure 13c,d). We set false discovery rate (FDR) ≤0.01 for both cell culture and kidney tissue RNASeq and added a threshold of fold change >2.0 for the tissue RNASeq with Anks^KO^. This yielded totals of 3250 and 4766 unique DEG combined from all genotype comparisons for kidney tissues and primary cells, respectively. In primary kidney cells, Pkd1^KO^ compared to WT had ∼3 fold higher number of DEG than Anks3^KO^ compared to WT (2833 vs. 911, respectively; Supplementary Figure 13c). In kidney tissue, Anks3^KO^ compared to WT had >20-fold higher DEG than Pkd1^KO^ compared to WT (2482 vs. 110, respectively). In keeping with the presymptomatic stage of Pkd1^KO^ at 7 weeks, both Pkd1^KO^ compared to WT and Pkd1^KO^+Anks3^KO^ compared to Anks3^KO^ alone had <200 DEG in kidney tissue that met the above criteria (Supplementary Figure 13d). In kidney tissues, Anks3^KO^, whether alone or in combination with Pkd1^KO^, had nearly 3-fold more upregulated than downregulated DEG, possibly reflecting expansion of new cell types in the kidney (Supplementary Figure13d-f). Pkd1^KO^ alone in kidney tissue and all DEG comparisons in primary cells had comparable numbers of up- and downregulated DEG (Supplementary Figure13c,d,g,h).

Unsupervised hierarchical clustering of the total unique DEG across all the pairwise comparisons of the kidney tissue genotypes showed that Anks3^KO^ and Pkd1^KO^+Anks3^KO^ had very similar patterns and were distinct from Pkd1^KO^ and WT, which in turn were similar to each other (Figure 6a). Genes that are most significantly altered in Anks3^KO^ compared to WT (*Serpina10*, *Lcn2*, *Lgals3*, *Timp1*, and *Tnfrsf12a*) and that had the largest absolute fold change (*Havcr1*, *Mmp7*, *Il1f6*, *Spink6*, *Trdn*) were also among the most significant changes or largest absolute fold changes, respectively, in Pkd1^KO^+Anks3^KO^ compared to WT (Supplementary Figure 13e,f). These included the kidney injury markers Ngal (*Lcn2*) and Kim1 (*Havcr1*). The predominance of the Anks3^KO^ genotype and limited impact of Pkd1^KO^ in kidney tissues is supported by gene ontology analyses of DEG showing similarly enriched pathways in Anks3^KO^ vs. WT and Pkd1^KO^+Anks3^KO^ vs. WT (Figure 6b,c). In both comparisons, regulation of leukocytes and immune response was strongly enriched in biological process (BP), organization of extracellular matrix was prominent in cellular component (CC), and cytokine-receptor activities were significantly enriched in molecular function (MF) (Figure 6b,c). These transcriptional responses correlate with inflammatory and fibrotic changes (Figure 5d-h) present in histologically normal appearing (Figure 4a,b) Anks3^KO^ kidneys at 7 weeks. By contrast, gene ontology analysis of DEG in Pkd1^KO^ tissues compared to WT showed no significant outcomes given DEG were <200. To increase the number of genes and ensure that we are not missing information, we performed gene ontology analysis using the Pkd1^KO^ vs. WT gene list defined by FDR ≤0.01 only (Supplementary Figure 13d). Even with this, Pkd1^KO^ vs. WT showed milder enrichment of more disparate pathways (Figure 6d) that differed from the changes in kidneys with Anks3^KO^.

**Figure 6.**
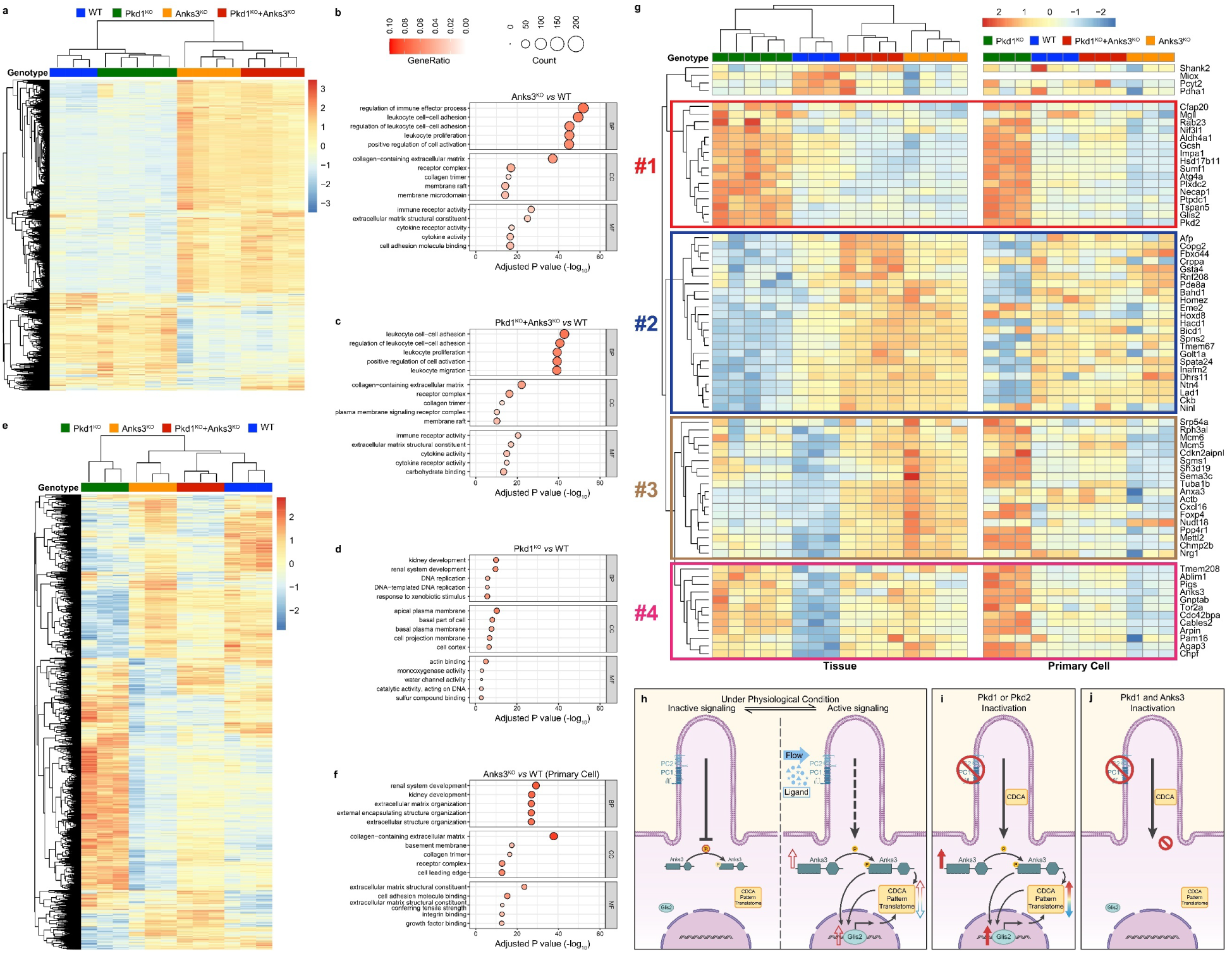
Anks3 has separate roles in CDCA signaling and kidney homeostasis. **a,e**, Heatmap showing unsupervised hierarchical clustering using default Euclidean distance metric for the 3250 (**a**) and 4766 (**e**) unique DEGs across all genotype comparisons from kidney tissues (**a**) and primary cells (**e**), respectively. Color scale indicates the relative gene expression and clustering by genotype is shown. **b-d,f**, Gene ontology analysis performed using the *clusterProfiler* R package with gene list in the indicated pairwise comparisons. Gene lists for Anks3^KO^ vs. WT (**b**), Pkd1^KO^+Anks3^KO^ vs. WT (**c**) in tissue were selected with FDR ≤0.01 and absolute two-fold change threshold. Gene lists for Pkd1^KO^ vs. WT (**d**) in tissue and for Anks3^KO^ vs. WT in primary cells (**f**) were selected only with FDR ≤0.01 threshold. BP, biological process; CC, cellular component; MF, molecular function. **g**, Heatmap showing unsupervised hierarchical clustering for the 73 core CDCA signature genes with the indicated genotypes in tissue RNASeq (*left panel*) and heatmap with the same order of genotypes and genes showing the 73 core CDCA signature genes in primary cell RNASeq (*right panel*). Color scale indicates the relative gene expression. Four clusters of gene (#1 — #4) are discussed in the text. **h-j**, A hypothesis for the role of Anks3 in CDCA. **h**, Under physiological conditions, PCs function as brakes that balance inactive (*left*) and active (*right*) signaling responses to flow or ligands to control tubule cell morphology. In the resting state (*left*), CDCA is quiescent and there is no upregulation of Anks3 and nuclear Glis2 and no change in the CDCA pattern translatome or basal phosphorylation of Anks3. In the active state (*right*), there are regulated increases in Anks3 and Glis2 expression, and changes in the CDCA pattern translatome and Anks3 phosphorylation. Glis2 upregulation in active signaling is dependent on Anks3. **i**, Inactivation of PCs causes constitutive CDCA signaling in primary cilia with persistent upregulation of Anks3 and consequent increase in nuclear Glis2 and activation of the CDCA pattern translatome. There is also altered Anks3 phosphorylation, although the direct role of that in CDCA is unknown. **j**, Inactivation of both Pkd1 and Anks3 does not alter the persistent CDCA signal in cilia, but the process is interrupted due to absence of Anks3 and there is no upregulation or nuclear expression of Glis2 and no changes in the CDCA pattern translatome.

In contrast to the tissue results, unsupervised hierarchical clustering of unique DEG in the primary cells had Pkd1^KO^ with the most distinct expression pattern (Figure 6e). The pattern of DEG in Anks3^KO^ alone showed an inverse effect to Pkd1^KO^, such that many genes upregulated in Pkd1^KO^ compared to WT were relatively downregulated in Anks3^KO^; genes downregulated in Pkd1^KO^ compared to WT were either unchanged or upregulated in Anks3^KO^ (Figure 6e). The Pkd1^KO^ DEG pattern in the primary cells reverted toward the WT expression pattern when combined with Anks3^KO^ in Pkd1^KO^+Anks3^KO^, showing that Anks3^KO^ has a broad suppressive effect on Pkd1^KO^ dependent gene expression changes (Figure 6e). These results support the idea that, at least for Anks3^KO^, the in vitro primary cell-based system is more reflective of cell autonomous changes, while the transcriptional response in bulk kidney tissue represents secondary, non-cell autonomous responses due to disrupted kidney homeostasis. Gene ontology analyses using primary cell DEG comparing Anks3^KO^ to WT (Figure 6f) showed enrichment of extracellular matrix related terms in all three categories (BP, CC, MF). DEG in Pkd1^KO^ in cells showed more disparate pathways without a consistent theme in BP, CC or MP (Supplementary Figure 13i). The absence of overlap in the early stage in vivo transcriptional responses between Anks3^KO^ and Pkd1^KO^ and the commonality of the changes observed in Anks3^KO^ and Pkd1^KO^+Anks3^KO^ support the conclusion that the Anks3^KO^ phenotype in the kidney is dominant and independent of PC1. However, in cells, *Anks3* inactivation has broad effect on transcriptional changes resulting from inactivation of *Pkd1*.

We considered that the interaction between *Anks3* and *Pkd1* cell autonomous transcriptional responses in vivo may be masked by the dominant kidney injury response to Anks3^KO^. To explore this directly, we focused specifically on the expression of the 73 CDCA pattern genes that are validated in vivo Pkd1^KO^ dependent changes (41). We performed unsupervised hierarchical clustering for the expression of these 73 CDCA pattern genes in kidney tissue samples for the four genotypes (Figure 6g, *left*). We then generated the heatmap for primary kidney cell RNASeq prespecifying the order of genotypes and genes determined by the hierarchical clustering in tissue (Figure 6g, *right*). The results can be grouped into four major clusters of genes (Figure 6g). Cluster #1 contains 16 genes, including *Glis2* and *Pkd2*, whose expression is upregulated in Pkd1^KO^ compared to WT and is normalized to even slightly below WT level in Pkd1^KO^+Anks3^KO^ in both kidney tissue and primary cells. Cluster #2 contains 23 genes, including *Spns2*, *Ntn4*, and *Lad1*, whose expression is downregulated in Pkd1^KO^ and is normalized to even slightly above WT level in Pkd1^KO^+Anks3^KO^ in both tissue and primary cells. The 18 genes in Cluster #3 follow the same pattern as in Cluster #1 in primary cells but not in tissue. The significant gene expression changes due to kidney injury may be obscuring the *Pkd1* dependent changes in these genes. The 12 genes in Cluster #4 again follow the same Pkd1^KO^ pattern as in Cluster #1 in primary cells and tissues, but the normalization by Pkd1^KO^+Anks3^KO^ is mild and intermediate in tissue. In aggregate, 48 of 73 CDCA pattern genes in primary kidney cells are significantly changed in the same direction in Pkd1^KO^ vs WT as we found in the TRAP RNASeq studies (41). Expression changes in 44 out of these 48 genes are suppressed toward WT levels in Pkd1^KO^+Anks3^KO^ cells. In tissue, 53 of 73 CDCA pattern genes are significantly changed in Pkd1^KO^ vs WT and changes in 46 out of the 53 are suppressed in Pkd1^KO^+Anks3^KO^. The suppressive effects by concomitant Anks3 inactivation are highly significant with *P*<2.2×10^-16^ by Fisher’s exact test in both cells and tissues. Overall, the data show that, in addition to suppressing Glis2 nuclear activation downstream of Pkd1^KO^, Anks3 inactivation suppresses expression changes in the majority of CDCA pattern genes both in vitro and in vivo (Figure 6h-j). The study defines Anks3 as a central cytoplasmic component of the CDCA signaling pathway acting downstream of polycystins and cilia and upstream of the CDCA effector Glis2.

## Discussion

The cilia dependent cyst activating (CDCA) signal whereby removal of cilia suppresses kidney tubule and bile duct cyst formation despite inactivation of polycystins (40, 52) defines a functional relationship between polycystins and primary cilia. CDCA behaves as a gain of function phenotype downstream of polycystin inactivation with intact cilia. CDCA chronically redefines tubule cell structure and function which in turn remodels the kidney leading to polycystic kidney disease with progressive kidney dysfunction. The molecular components of the cellular pathways regulated by CDCA were unknown. To address this gap in understanding, we developed an unbiased in vivo discovery platform using translating ribosome affinity purification (TRAP) RNASeq in tripartite comparisons between control, Pkd^KO^, and Pkd^KO^+cilia^KO^ mutant mouse kidneys to define a tubule cell-specific CDCA pattern translatome in precystic kidneys (41). From this we previously identified *Glis2*, encoding a nuclear transcription factor, as a functional effector of polycystin and CDCA signaling and showed that Glis2 upregulation in kidney tissues and cell cultures is a robust readout for polycystin-dependent cyst forming signals (41). This left a gap in understanding of how a cilia based signal, i.e., CDCA, results in upregulation at the transcriptional or post-transcriptional level of Glis2 mRNA. To address this gap, we applied a new paradigm for discovery of additional components of CDCA by assaying for suppression of Glis2 upregulation in polycystin knockout cells. While such candidate CDCA signaling components are not a priori expected to show CDCA pattern transcriptional changes, we nonetheless hypothesized that the CDCA translatome would be a good place to begin to look for candidates. We found that inactivation of *Anks3*, which shared a CDCA pattern expression profile very similar to *Glis2* (41), can completely suppress *Glis2* upregulation at both the mRNA and protein level in Pkd1^KO^ with intact cilia. Given the very significant suppression of Glis2 upregulation by Anks3 inactivation and the knowledge that Glis2 upregulation appears to be necessary for cyst growth (41), we hypothesized that Anks3 is a central element of the CDCA pathway.

Anks3 is an attractive candidate for a functional mediator of CDCA given that it is an interacting protein regulator for nephronophthisis (NPHP) and cilia-related gene products (47, 48, 50, 51, 53, 54). The structural features (ankyrin and SAM domains) (55, 56) and interaction profiles suggests Anks3 functions as a scaffolding protein for integrating cilia-related signaling (49, 54, 57, 58). Given these interactions, we considered whether the role of Anks3 in CDCA may be upstream of cilia, controlling cilia composition in a manner analogous to Tulp3 (59, 60). Several data, however, suggest that Anks3 function is downstream of polycystins and cilia. First, Anks3 is upregulated by Pkd1^KO^ and this upregulation is suppressed by polycystin and cilia dual inactivation, i.e., the CDCA pattern, so it appears regulated by both polycystins and cilia making it more likely that it functions downstream of both. Second, the kidney injury that occurs following Anks3^KO^ is not impacted by cilia inactivation showing that the role of Anks3 in kidney homeostasis is not cilia dependent. Third, Anks3 is a cytosolic protein that does not enter cilia. The most direct interpretation of these results is that Anks3 is a CDCA component downstream of polycystin function in cilia. The placement of Anks3 upstream of Glis2 in CDCA is more definitive. Anks3 inactivation suppresses Glis2 very efficiently but Glis2 inactivation has no effect on Anks3 upregulation or phosphorylation state.

The role Anks3 as a scaffolding protein may provide clues to its possible functions in CDCA. It can be proposed that Anks3 has quantitative and qualitative changes in scaffolding function of other bioactive signaling components under control of CDCA. In this regard, we found that polycystin function regulates the phosphorylation state of Anks3. This in turn may affect its interactome to alter signaling under Pkd1^KO^ conditions [e.g., (61)]. Anks3 also has cytosolic interactors such as Bicc1, which has been implicated in modulation of mRNAs related to ciliopathy phenotypes (47, 51, 58, 62), so there is basis for a hypothesis that this may play a role in regulating steady state mRNA levels of CDCA components including *Glis2*. In addition to its role in CDCA, we found that Anks3 has a profound impact on kidney tubule structural and functional homeostasis. The role of Anks3 in maintaining the integrity of the adult kidney was not previously appreciated. The effects are profound and rapid compared to the timescale of ADPKD in adult animal model, indicating that Anks3 has an active role with a short timescale in maintaining the kidneys’ structure. The primary kidney cell transcriptomic studies coalesce around a significant role for Anks3 in maintaining extracellular matrix (ECM) while the whole kidney transcriptomic data suggest that disruption of this function in vivo promotes an aggressive immune mediated response. Future studies should focus on the specific ECM responses that depend on Anks3. These can define specific elements of kidney ECM that, when disrupted, promote immune damage to tubules.

While these mechanisms define a broader role for Anks3 in the kidney beyond functional dependence on polycystins and cilia, the most striking outcome regarding ADPKD is the profound extent to which Anks3 impacts the CDCA translatome specifically. This is seen in both primary cell culture models and in kidney tissues; the latter despite the profound non-CDCA related effects Anks3 has on the kidney. Together these findings indicate a central role for Anks3 in CDCA. The suppression of so many of the CDCA transcriptional changes in both cell and kidney models by inactivation of Anks3 leaves open the possibility that Anks3 is the cytosolic nidus of control for CDCA. There are two broad interpretations of the suppression of CDCA pattern transcriptional changes by Anks3^KO^; the first is that it is largely mediated by Anks3 through suppression of Glis2 and the other is that Anks3 regulates multiple arms of cellular CDCA signaling, of which Glis2 is one. Of these, the former is the more attractive hypothesis given its simplicity. These two hypotheses can be resolved once the functional relationship of Glis2 transcription factor activity to the CDCA pattern translatome is defined—does Glis2 directly regulate expression of the genes in the CDCA translatome at the level of transcription? If Anks3 exerts control of CDCA primarily through Glis2, then the focus will be to determine how Anks3 regulates Glis2 mRNA steady state levels. The studies presented here provide progress in understanding polycystin and cilia dependent pathways that result in polycystic kidney disease following loss of polycystins. ADPKD results from increased activity of Anks3 which results in increased activity of the transcription factor Glis2. Glis2 remains a viable therapeutic target (41) while Anks3 has other critical roles in the adult kidney that make it a less attractive direct target for therapy. Evidence suggests that the role of Anks3 in polycystic kidney disease is functionally distinct from its role in adult kidney homeostasis. Identifying the function of Anks3 specifically in CDCA driving ADPKD will advance understanding of the cell autonomous molecular bases of ADPKD and may identify additional therapeutic targets that are specific and effective for ADPKD.

## Methods

### Mouse strains and procedures

All experiments were conducted in strict accordance with the guidelines and procedures set by the Institutional Animal Care and Use Committee (IACUC) at Yale University. The mice used are >95% congenic on the C57BL/6 background. Mice of both sexes were used unless otherwise indicated. The following strains of mice used in this study were previously described: *Pkd1^fl^* (63), *Pax8^rtTA^* (JAX Strain # 007176), *TetO^Cre^* (JAX Strain # 006234), *Ift88^fl^* (JAX Strain # 022409) (64), *Pkd2^fl^* (65), *Pkd2^-^* (null) (9), *Pkd2^FSF^-BAC* (66), and *Rosa^FlopER^* (66). The *Glis2^HT^* is briefly described (Dong et al., manuscript in preparation). The *Glis2^HT^* allele was generated using CRISPR/Cas9 to target the C-terminus of endogenous *Glis2* allele. A single-stranded oligodeoxynucleotides (ssODNs) containing the *HaloTag7* sequence with two homology arms was designed as repair template for precise genome editing via homologous recombination. Cas9 protein, mouse *Glis2* guide RNA and ssODNs were microinjected into the fertilized embryos and then transferred into pseudo-pregnant wildtype mothers to generate knock-in founder lines which were produced in the Yale Genome Editing Center in (C57BL/6J X SJL/J) F2 zygotes. Founders were identified by PCR genotyping and verified by sequencing of PCR products as well as immunoblotting of the fusion protein. *Anks3* conditional and null allele were produced in this study by the Yale Genome Editing Center through CRISPR/Cas9 targeting. Detailed description provided in supplementary information.

Gene inactivation in the early-onset models with the *Pax8^rtTA^; TetO^Cre^* digenic system was induced with 2 mg/ml of doxycycline in drinking water supplemented with 3% sucrose provided to the nursing dams for two weeks from postnatal day 0 (P0) to P14. All pups were examined at P14. Gene inactivation in the adult-onset models of *Pax8^rtTA^; TetO^Cre^*was induced with 2 mg/ml of doxycycline in drinking water supplemented with 3% sucrose given to mice for two weeks from P28 to P42. Adult model mice were euthanized at 7, 10, 13, or 19 weeks of age. Genotyping was done on DNA isolated from toe clips. Copy number of Pax8^rtTA^ and TetO^Cre^ was determined as previously reported (67). Blood samples were obtained by ventricular puncture. Serum was separated using Plasma Separator Tubes with lithium heparin (BD Biosciences, Cat. no. 365985). Serum urea nitrogen was analyzed by the George M. O’Brien Kidney Center at Yale University. One kidney was snap-frozen for protein and mRNA extraction, and the other kidney was fixed in 4% paraformaldehyde for histological analysis. Sagittal sections of kidneys were processed for hematoxylin and eosin (H&E), periodic acid-Schiff (PAS), and Masson-trichrome staining. Cystic index was calculated as previously described (63).

### Primary cell culture

Primary cells from mouse kidneys were isolated and cultured as previously described (67). Cells were let to attach for 48-72 hours (h) and then treated with doxycycline (Sigma, Cat. no. D9891-100G) 1 µg/ml for 72 h. For selected experiments, cells were further treated with 4-hydroxy Tamoxifen (Cayman Chemical, Cat. no.17308) at 4 µM for 72 h. All cells were serum starved to promote cilia formation in 0.1% FBS containing media for 24 h prior to preparation of protein or RNA.

### cDNA constructs, transfection, and cell culture

The full-length cDNA of mouse *Anks3* (NCBI Accession NM_028301.5) was cloned into several vectors: pLX304 (Addgene, Cat. no. 25890) with C-terminal triple HA tag; pcDNA 3.1(-)/myc-His C (Thermo fisher V85520) with both N-terminal V5 and C-terminal triple HA tag; and C-terminal EGFP tagged Anks3 was cloned with the cilia marker Nphp3^(1-200)^-mApple (67) with a tandem P2A-T2A (tPT2A) linker (68, 69) for bi-cistronic gene expression into pcDNA 3.1(-)/myc-His C. Generation of immortalized cell lines from primary cells using lentivirus of SV40 large T antigen (Gentarget, Cat. no. LVP016-Hygro) was performed as previously described (67). Immortalized cell lines and IMCD3 cells (ATCC CRL-2123) were transfected with plasmid DNA using Lipofectamine LTX with Plus reagent (Invitrogen, Cat. no. 15338100) and selected with Blasticidin (10 µg/ml, InvivoGen, Cat. no. ant-bl-1) or G418 (500 µg/ml, Sigma-Aldrich, Cat. no. G8168-100ML) 48 h after transfection for stable expression. A Pkd1^KO^ LLC-PK_1_ cell line was generated by CRISPR/Cas9 targeting the exon 2 and 3 of *Pkd1* gene, followed by clonal isolation. Wild type LLC-PK_1_ cells and the L2-C2 *Pkd1*^KO^ LLC-PK_1_ clone were transfected with plasmid DNA through electroporation and were selected with G418 (500 µg/ml) 48 h after electroporation. All cell lines were cultured in DMEM high glucose (Thermo Fisher Scientific, Cat. no. 11965092) with 5% FBS and were serum starved in 0.1% FBS containing media for 24 h prior experimentation.

### In vitro gene knockout

*In vitro* gene knockout studies were done in immortalized cell lines or primary cells using CRISPR/Cas9 and nucleofection. Ribonucleoprotein (RNP) complexes composed of 1.5 µl of NLS-Cas9-EGFP Nuclease (Genscript, Cat. no. Z03393-100), 3 µl of reconstituted sgRNA pool (three sgRNAs per gene, ordered from Synthego), and 2 µl of 1X TE buffer were prepared freshly and incubated at room temperature for 15 min. The RNP complexes were then mixed with 100,000 cells resuspended in 20 µl of nucleofector solution. The mixture was nucleofected with 4D-Nucleofector X Unit (Lonza) and program CM-137 in 16-well format plate and then resuspended and transferred to multiwell plates and allowed to grow until confluent. For selected experiments, single-cell clones were isolated by serial dilution, and verified by Sanger sequencing and immunoblotting. The following gRNAs were used:

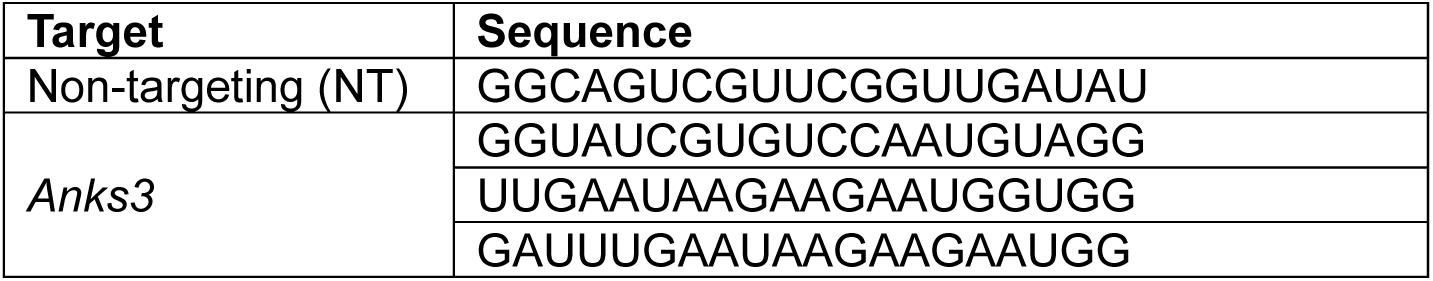

### Immunocytochemistry

Immunofluorescence cell staining was performed on immortalized cell lines and primary cells with standard protocols. For immortalized cells with Glis2^Halo^, cells were incubated with HaloTag TMR ligand (1:10000 dilution in media, Promega, Cat. no. G8251) for 30 min at 37°C before fixation. All adhered cells were fixed with cold methanol for 15 min at 4°C, followed by blocking in 3% BSA in PBS for 30 min at room temperature. Primary antibodies were diluted in 3% BSA and incubated overnight at 4°C. Cells were washed 3 times with 0.1% Tween-20 in PBS (PBST) followed by secondary antibody in 3% BSA for 1 h at room temperature. LLC-PK_1_ cells were imaged as non-fixed and live cells. Immunofluorescence of kidney cryosections (5-7 µm) was performed according to standard procedures as well. Briefly, sections were permeabilized with 0.2% Triton X-100 in PBS for 30 min and blocked with 0.1% BSA and 10% goat serum in PBS for 1 h at room temperature, followed by primary antibody incubation overnight at 4°C and secondary antibody incubation for 1 h at room temperature. All images were acquired on Nikon Eclipse Ti (Nikon Instruments Inc, Japan) equipped with Yokogawa CSU-W1 spinning disc and Andor solid state lasers (Andor Technology, UK), using either NIS-Elements AR (Nikon, Version 4.30.02) or MetaMorph imaging software (Molecular Devices LLC, Version 7.10.5.476). The following antibodies and lectins were used: rat anti-HA (1:200, Roche, Cat. no. 11867423001); rhodamine *Dolichos biflorus agglutinin* (DBA) (1:200, Vectors Laboratories, Cat. no. RL-1032); FITC *Lotus tetragonolobus agglutinin* (LTA) (1:200, Vectors Laboratories, Cat. no. FL-1321); rabbit anti-megalin (1:1000) (70); rabbit anti-Ki67 (1:200, Cell Signaling Technology, Cat. no. 12202S); rat anti-F4/80 (1:200, AbD Serotec, Cat. no. MCA497R); mouse anti-TNFα (1:200, Santa Cruz Biotechnology, Cat. no. SC-52746); rabbit anti-PDGFRβ (1:200, Abcam, Cat. no. ab32570); rabbit anti-αSMA (1:200, Abcam, Cat. no. ab5694); mouse anti-Arl13b (1:1000, Proteintech, Cat. no. 66739-1-lg); rabbit anti-Anks6 (1:100, Sigma Aldrich, Cat. no. HPA008355); rabbit anti-Invs (1:100, Proteintech, Cat. no. 10585-1-AP); rabbit anti-PC2 (YCC2 (71), 1:1000); and Alexa-488/597/647-conjugated secondary antibodies (1:200, Molecular Probes). Hoechst 33342 (1:5000, Molecular Probes, Cat. no. H3570) was used for nuclei staining.

### Generation of anti-Anks3 antibodies

Anti-Anks3 antibodies were custom produced by Covance (now Labcorp Drug Development, NC, USA). Two rabbits were injected with 500 µg of each of two peptides corresponding to amino acids 384-399 and 615-629 of the mouse Anks3 sequence (NCBI Accession NP_082577.2) followed by three rounds of boosting. Serum was collected after final boost and antibodies were obtained from the pooled serum by peptide antigen affinity purification.

### Protein preparation, immunoblotting, and immunoprecipitation

Total cell lysates were prepared with 1x Red Loading Buffer Pack (Cell Signal Technology, Cat. no. 7723S) as previously described (67) or with ice-cold lysis buffer (50 mM Tris-HCl pH 7.5, 150 mM NaCl, and 1% NP-40) supplemented with complete EDTA-free protease inhibitor cocktail tablets (Roche, Cat. no. 11836170001) and PhosSTOP phosphatase inhibitor cocktail tablets (Roche, Cat. no. 4906845001). Total cell lysates used for phosphatase experiment were prepared similarly but without PhosSTOP phosphatase inhibitor and were treated with Lambda Protein Phosphatase (NEB, Cat. no. P0753S) for 30 min at 30°C based on manufacturer’s instructions. Total kidney lysates were prepared in ice-cold 1X RIPA buffer (Abcam, Cat. no. ab156034) with the same supplements and homogenized in a bead mill homogenizer (Precellys^®^ Evolution, Bertin Instruments) at 6800 rpm for 30 s twice with 1 min ice-bath in between. Lysates were centrifuged at max speed for 10 min at 4°C. Fractionation of cells or kidney tissues into cytosolic and nuclear fractions were done using the NE-PER^TM^ Nuclear and Cytoplasmic Extraction Reagents (Thermo, Cat. no. 78833) with manufacturer’s instructions. Protein concentrations were measured with bicinchoninic acid assay (BCA) kit (Thermo Fisher Scientific, Cat. no. 23225) or Protein Assay Dye Reagent Concentrate (Bio-Rad, Cat. no. 5000006). Equal amounts of total protein were loaded and separated in 4-20% Mini-Protean TGX Precast Gels (Bio-Rad, Cat. no. 4568094) and transferred to Nitrocellulose membrane (Bio-Rad, Cat. no. 1620115). Membranes were blocked with 5% milk for 1 h, incubated with primary antibodies overnight at 4°C, and with secondary antibodies for 1 h at room temperature. Analysis of phosphorylated proteins using 7.5% SuperSep Phos-tag (50 µmol/L) Precast Gels (Wako Chemicals, Cat. no. 198-17981) was performed according to manufacturer’s instructions. Membrane stripping and re-probing were done using Restore PLUS Western Blot Stripping Buffer (Thermo Scientific, Cat. no. 46430). The images were acquired with LI-COR Odyssey Fc Imaging system, and the densitometric quantifications of immunoblot bands were analyzed with Image Studio Lite software (LI-COR Biosciences, Version 5.2.5).

The following primary antibodies were used: rabbit anti-Anks3 (1:1000); rabbit anti-Glis2 (YNG2, (67) 1:20000); rabbit anti-Hsp90 (1:1000, Cell Signaling Technology, Cat. no. 4877S), mouse anti-HaloTag (1:1000, Promega, Cat. no. G9211); rat anti-HA (1:1000, Roche, Cat. no. 11867423001); rabbit anti-Lamin B1 (1:1000, Proteintech, Cat. no. 12987-1-AP); rabbit anti-Lamin A/C (1:1000, Cell Signaling Technology, Cat. no. 2032S); rabbit anti-phospho-Akt (Ser473) (1:1000, Cell Signaling Technology, Cat. no. 4060T); mouse anti-V5 (1:1000, Thermo Scientific, Cat. no. R960-25). Secondary anti-rabbit horseradish peroxidase (HRP), anti-mouse HRP and anti-rat HRP (1:5000, Jackson ImmunoResearch) were used.

### Mass spectrometry and proteomics

Immunoprecipitation of Anks3 from total cell lysates of primary cultures of *Pkd1^fl/fl^; Pax8^rtTA^; TetO^Cre^* mice (n=3), with (Pkd1^KO^) or without (WT) in vitro doxycycline induction, was done by mixing lysates with anti-Anks3 (1:25) and Pierce Protein G Magnetic Beads (Thermo Scientific, Cat. no. 88848) and incubating with rotation overnight at 4°C. Beads were washed five times with wash buffer (50 mM Tris-HCl pH 7.5, 150 mM NaCl, and 0.1% NP-40), and samples were eluted in 1X loading buffer (National Diagnostics, Cat. no. EC-887) by boiling. PAGE gels were run as above and protein bands were visualized using SimplyBlue SafeStain (Thermo Scientific, Cat. no. LC6060). One small gel band corresponding to 75 kDa was excised per sample and stored at -80°C until proceeding to mass spectrometric analysis.

LC-MS/MS was performed by the Keck MS & Proteomics Resource at Yale School of Medicine. A label free quantitation (LFQ) approach utilizing Progenesis QI (Nonlinear Dynamics, version 4.2) similar to that found in Vidyadhara et al (72) was utilized to obtain quantitative information on peptides and proteins. Briefly, the Progenesis QI software performs chromatographic/spectral alignment, mass spectral peak picking and filtering, and quantitation of proteins and peptides. A normalization factor for each run was calculated to account for differences in sample load between injections as well as differences in ionization. The algorithm then calculates the tabulated raw and normalized abundances and ANOVA P values for each feature in the data set. The MS/MS spectra was exported as .mgf (Mascot generic files) for database searching. Mascot Distiller was used to generate peak lists, and the Mascot search algorithm was used for searching against the Swiss Protein database with taxonomy restricted to *Mus musculus*; and carbamidomethyl (Cys), oxidation of Met, Phospho (Ser, Thr, Tyr), acetylation (Lys and Protein N-term), and deamidation (Asn and Asp) were entered as variable modifications. The Mascot search results were exported as .xml files and then imported into the processed dataset in Progenesis QI software where peptides ID’ed were synched with the corresponding quantified features and their corresponding abundances.

*RNA isolation, RT-qPCR, and RNASeq analysis.* Total RNA from cells or whole kidneys were isolated using Trizol (Themo Fisher Scientific, Cat. no. 15596026) and RNeasy Mini Kit (Qiagen, Cat. no. 74104) and used for cDNA synthesis by iScript cDNA synthesis kit (Bio-Red, Cat. no. 1708890) or used for RNA sequencing directly. RT-qPCR was performed using iTaq Universal SYBR green Supermix (Bio-Rad, Cat. no. 18064022) in CFX96 Touch Real-Time PCR detection system (Bio-Rad, USA). The following primers for RT-qPCR were used:

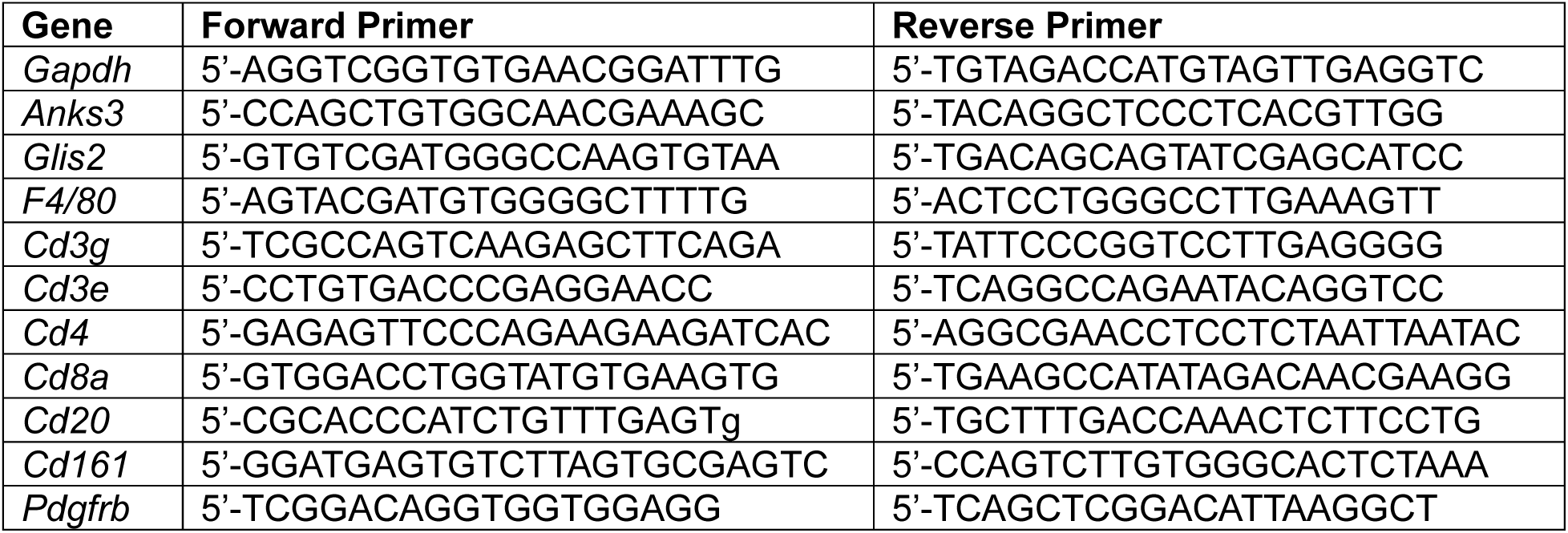

RNA sequencing library was prepared using the ribosomal depletion method (KAPA RNA HyperPrep Kit with RiboErase, Roche). Sequencing was run on Illumina NovaSeq 6000 platform using 150 bp paired end reads with read depth 40 or 100 million reads per sample, for cells or tissue respectively. RNA sequencing analysis for raw reads processing, cleaned reads alignment, and expression quantification of aligned reads were performed based on previously published pipeline (67). We only focused on protein coding genes. The filtered read counts matrix was normalized by the transcripts per million (TPM) method. Detection of differentially expressed genes was performed using R package *DESeq2* (73) and the Benjamini–Hochberg procedure was used for multiple test correction with FDR ≤0.01 alone or FDR ≤0.01 with absolute two-fold change cut-off used as the significance threshold for detection of differentially expressed genes (DEGs) in primary cells and kidney tissues, respectively. Heatmaps were generated using the *pheatmap* package in R with input matrix consisted of log_2_(TPM+1) transformed gene expression values. Both rows (genes) and columns (samples) were subjected to hierarchical clustering using default Euclidean distance metric to visualize patterns of expression across samples. Gene ontology analyses were performed using the *clusterProfiler* (74) R package.

### Statistics

Most of the quantitative data were analyzed using one-way analysis of variance (ANOVA) followed by Tukey’s multiple-comparison test. The rest of quantitative data were analyzed using two-tailed, unpaired Student’s *t* test as indicated in figure legends. All data are presented as mean ± s.e.m, and *P* < 0.05 was used as the threshold for statistical significance. GraphPad Prism (10.3.0) software was used to perform statistical analyses.

### Study approval

All animal studies were reviewed and approved by the IACUC of Yale University.

### Data availability

All the raw sequencing data and processed data have been deposited in the Gene Expression Omnibus (GEO) with the following accession number: GSEXXXXXX (pending). Access is not restricted as of the date of peer reviewed publication. The analytic methods followed the previous publication (67). The raw mass spectrometry/proteomics data have been deposited in the Proteomics Identifications Database (PRIDE) with the following accession number: PXDXXXXXX (pending) and are publicly available as of the date of publication.

## Supporting information

Combined file with supplementary information and supplementary figures

## Author Contributions

Z.W. designed and performed experiments, analyzed data, and drafted figures and manuscript. This work was performed as part of the fulfilment of doctoral thesis requirements for Z.W.

M.R. and K.R. performed selected experiments.

K.D., A.C., M.S.T. provided samples from re-expression mouse models.

K.D. generated and provided Glis2^Halo^ mouse line.

M.R., K.D., Y.C. and X.T. provided reagents.

J.G., supervised by H.Z., performed bioinformatic analyses.

G.L. performed pathology assessment and acquired histology images.

J.K. and T.L. from Yale Keck facility performed and analyzed LC-MS/MS data.

S.S. conceived the study, designed experiments, supervised the study, and wrote the manuscript.

## Acknowledgements

This work was supported by NIH/National Institute of Diabetes and Digestive and Kidney Diseases grants (nos. R01 DK120911, R01 DK100592, and RC2 DK120534 to S.S.) and a grant from the Amy P. Goldman Foundation to S.S. We are grateful for the generous support from Mr. and Mrs. Robert Roth. We also thank the Keck MS & Proteomics Resource at Yale School of Medicine for providing the necessary mass spectrometers and the accompany biotechnology tools funded in part by the Yale School of Medicine and by the Office of The Director, National Institutes of Health (S10OD02365101A1, S10OD019967, and S10OD018034). The funders had no role in study design, data collection and analysis, decision to publish, or preparation of the manuscript.

## Conflict of interest

The authors have declared that no conflict of interest exists with this work.

## Notes

### Competing Interest Statement

The authors have declared no competing interest.

