## Supplementary material for "Anks3 mediates cilia dependent polycystin signaling and is essential for adult kidney homeostasis": Combined file with supplementary information and supplementary figures

*Generation of a conditional Anks3$^{fl}$ mouse allele and a Anks3$^{\Delta 3\text{-}4}$ null allele*

We generated a Cre recombinase inducible conditional allele of *Anks3* (*Anks3$^{fl}$*) by inserting a pair of *lox*P sites flanking exons 3 and 4, which encode for the second to fourth ankyrin repeats (Supplementary Figure 3a). The founder mouse had one chromosome with the intended conditional allele (Supplementary Figure 3b), and the other chromosome with a deletion between the two guide RNA cleavage sites (*Anks3$^{\Delta 3\text{-}4}$*) (Supplementary Figure 3c). We separated the two alleles in the F1 generation. Intercrossing heterozygous *Anks3$^{\Delta 3\text{-}4/+}$* mice did not result in any viable postnatal homozygous *Anks3$^{\Delta 3\text{-}4/\Delta 3\text{-}4}$* mice (Supplementary Figure 3d). Embryos examined at E15.5 showed the expected Mendelian ratios (Supplementary Figure 3e) and evidence of abnormal left-right axis formation, consistent with a previous report showing embryonic or perinatal lethality with randomization of organ laterality (1). This shows that our deletion of exons 3 and 4 results in complete loss of function for *Anks3*. The conditional *Anks3$^{fl}$* allele was transmitted in normal Mendelian ratios and had no discernible abnormal phenotype in the homozygous state.

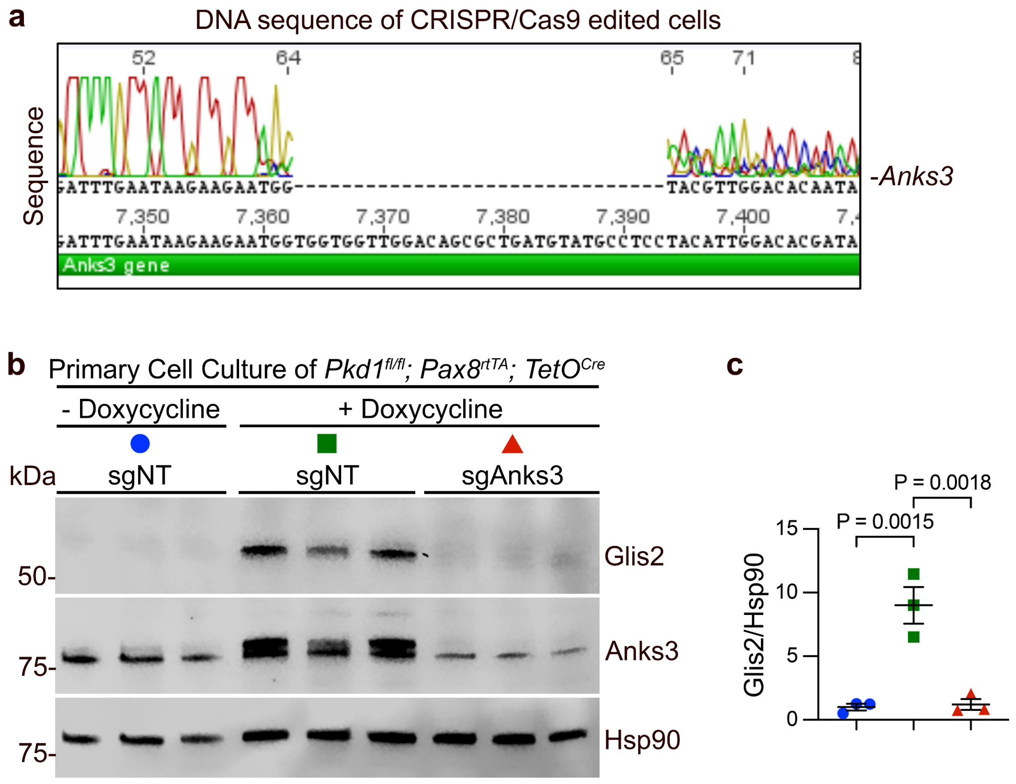


**Supplementary Figure 1. Acute CRISPR inactivation of Anks3 by nucleofection.** **a**, Sanger sequencing trace showing the success of Anks3 knockout. **b**, Immunoblots and **c**, quantification of whole cell lysates from renal primary cell cultures of *Pkd1*$^{fl/fl}$; *Pax8*$^{rtTA}$; *TetO*$^{Cre}$ mice with or without in vitro doxycycline treatment, followed by acute CRISPR/Cas9 gene targeting by non-targeting (sgNT) or Anks3 (sgAnks3) guide RNAs. Fold changes of the ratio of Glis2 to Hsp90 for each lane are shown relative to the mean of the "-doxycycline with sgNT" group, which is set to 1.0. Multiple-group comparisons were performed by one-way ANOVA followed by Tukey's multiple-comparison test, presented as mean ± s.e.m.

**a**

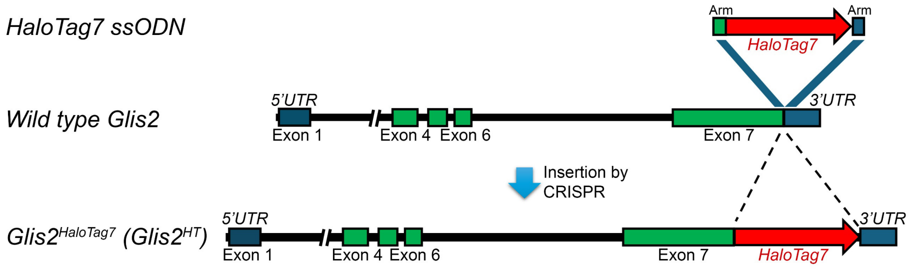


**b**

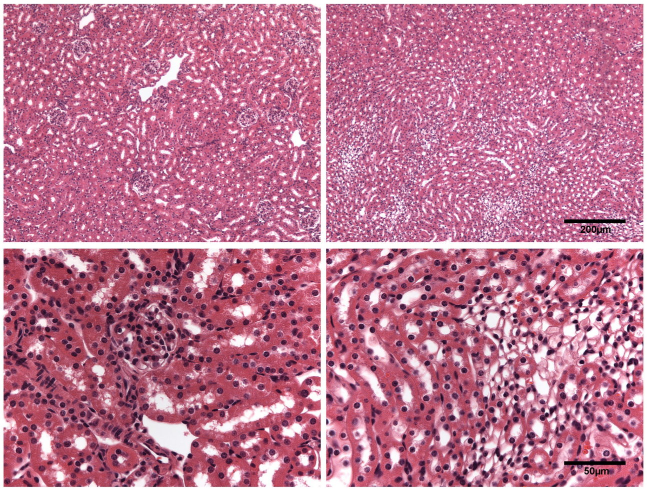


**Supplementary Figure 2. Generation of Glis2-HaloTag7 allele.** **a**, Strategy for generation of mouse *Glis2-HaloTag7* (*Glis2*$^{HT}$) knockin on chromosome 16. *HaloTag7* was inserted just after mouse exon 7 and before stop codon in endogenous *Glis2* in mouse. **b**, Representative hematoxylin and eosin stained histological kidney sections from cortex (left) and medulla (right) of 26-week old *Glis2*$^{HT/HT}$ (Glis2$^{Halo}$) mice showing normal renal histology without evidence of inflammation, fibrosis or other kidney damage. Scale bars, upper panels, 200 μm; lower panels, 50 μm.

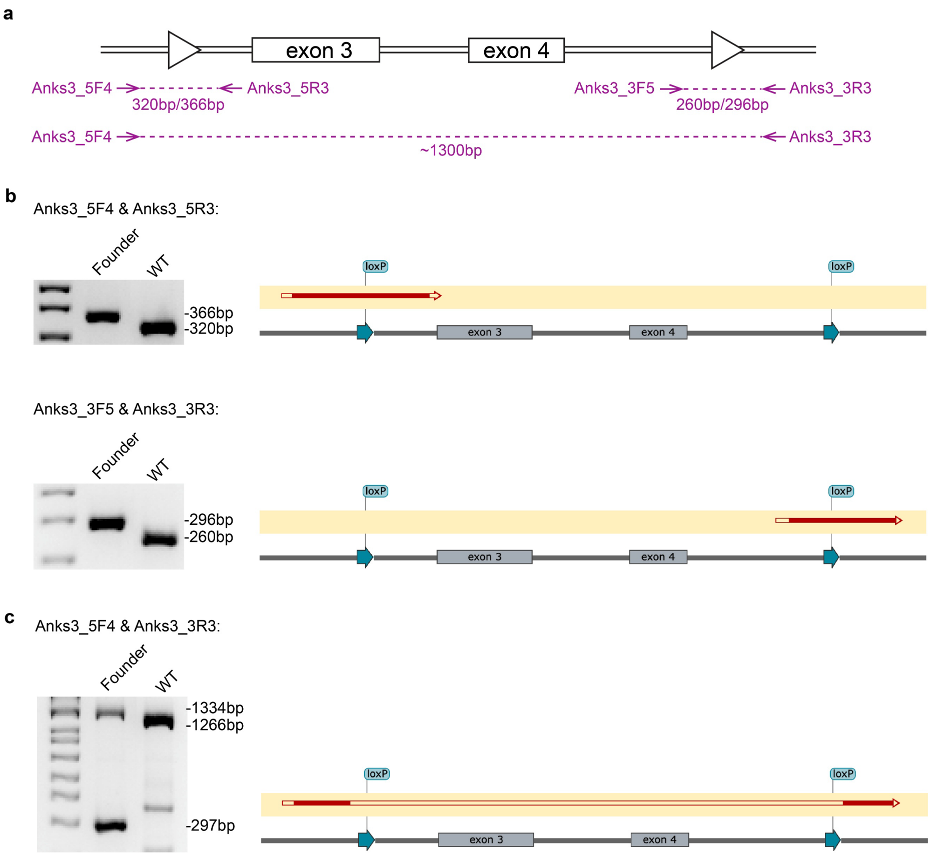


**d**

$Anks3^{+/\Delta 3\text{-}4}$ X $Anks3^{+/\Delta 3\text{-}4}$:

| | $Anks3^{+}$ | $Anks3^{\Delta 3\text{-}4}$ |
|---|---|---|
| $Anks3^{+}$ | $Anks3^{+/+}$ 82 (35.7%) | $Anks3^{+/\Delta 3\text{-}4}$ 148 (64.3%) |
| $Anks3^{\Delta 3\text{-}4}$ | $Anks3^{+/\Delta 3\text{-}4}$ 148 (64.3%) | $Anks3^{\Delta 3\text{-}4/\Delta 3\text{-}4}$ 0 (0%) |

**e**

$Anks3^{+/\Delta 3\text{-}4}$ X $Anks3^{+/\Delta 3\text{-}4}$ (E15.5):

| | $Anks3^{+}$ | $Anks3^{\Delta 3\text{-}4}$ |
|---|---|---|
| $Anks3^{+}$ | $Anks3^{+/+}$ 13 (22.8%) | $Anks3^{+/\Delta 3\text{-}4}$ 28 (49.1%) |
| $Anks3^{\Delta 3\text{-}4}$ | $Anks3^{+/\Delta 3\text{-}4}$ 28 (49.1%) | $Anks3^{\Delta 3\text{-}4/\Delta 3\text{-}4}$ 16 (28.1%) |

**Supplementary Figure 3. Generation of conditional *Anks3$^{fl}$* allele and *Anks3$^{\Delta 3\text{-}4}$* null allele.** **a**, Schematic of *Anks3* conditional allele (*Anks3$^{fl}$*) and the location of genotyping primers (purple arrows). **b**,**c**, Agarose gel images and sequencing results from founder mice to show one chromosome with the intended conditional allele (*Anks3$^{fl}$*, **b**), and the other chromosome with a deletion between the two guide RNA cleavage sites (*Anks3$^{\Delta 3\text{-}4}$*, **c**). Red filled arrow segments show matching sequences. **d**,**e**, Punnett square with numbers and percentages at postnatal (**d**) and E15.5 (**e**), showing the late embryonic lethality of Anks3 homozygous (*Anks3$^{\Delta 3\text{-}4/\Delta 3\text{-}4}$*) mice.

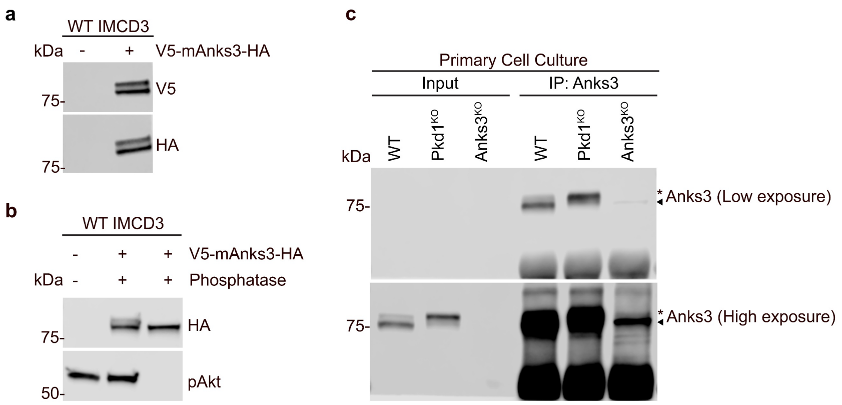


**Supplementary Figure 4. Anks3 is phosphorylated. a,b**, Heterologously expressed Anks3 is phosphorylated. **a**, Total cell lysate of wild type (WT) IMCD3 cells over-expressing N-terminal V5 and C-terminal triple HA tagged mouse Anks3 (V5-mAnks3-HA) immunoblotted with anti-V5 and anti-HA antibody showing two bands. **b**, Total cell lysate of WT IMCD3 cells over-expressing V5-mAnks3-HA was treated with Lambda protein phosphatase and immunoblotted with anti-HA showing disappearance of the upper band after phosphatase treatment. Phospho-Akt was used as positive control for phosphatase activity. **c**, Anti-Anks3 antibody can immunoprecipitate all phosphorylation states of Anks3. Primary kidney cell cultures from wild type, $Pkd1^{fl/fl}$*;* $Pax8^{rtTA}$*;* $TetO^{Cre}$ (Pkd1$^{KO}$), and $Anks3^{fl/fl}$*;* $Pax8^{rtTA}$*;* $TetO^{Cre}$ (Anks3$^{KO}$) mice were treated in vitro with doxycycline. Total cell lysates were immunoprecipitated with anti-Anks3 antibody, showing that all forms of Anks3 can be immunoprecipitated and enriched. High exposure of the blot was used to show the signal in input samples.

**a**

**WT:**

**Protein sequence coverage: 74%**

1 M**SELSDEASE PELLKRSLSM WHGLGAQGSP EELDVPLDLH TAASIGQHEV**
51 **VK**ECVQR**GEL DLNKK**NGGGW TALMYASYIG HDTIVHLLLE AGVSVNVPTP
101 EGQTPLMLAS SCGNESIAYF LLQQGAELEM KDIHGWTALF HCTSAGHQQM
151 VK**FLLESGAN ANVREPVYGY TPLMEAAASG HEIIVQYFLN HGVK**VDTR**DH**
201 **SGATACMLAR** QFGHMK**IVAL METHSPVLPK** SLYR**SPEKYE DLSSSDESWP**
251 **VPQR**QRPCRK **KGLSIHEGPR** ALAK**ITAIGL GGKTQTTYEQ VPPRGYVTFT**
301 **SSDENTMESE GLCYRDVTSP INEQDVESSS SSSREEPTFC ASLGPVWRSS**
351 **SSDGLARAQG LSSEASIESN EDSDHARK**SS VRKQTRSYLK NKSR**HGNSDG**
401 **HWPSSTGPAS IPGSEPQTEK SPYSGPQDLA TLLEQIGCLK YLQVFEEQDI**
451 **DLRIFLTLTE SDLKEIGITL FGPK**RKMTSA IAR**WHSSARP PSDALELAYA**
501 **DRLETEMQEL AIQLHK**CCEE AEALR**GQVSQ EQELRAVVES CLLEQDSAR**K
551 **DIHAQLQEAQ TLAQDAALVL DQLRACQAEL SAR**LRQHHSP R**EGTPNPHFL**
601 **SADSKGWPIP LQALSLPELS GALEDRVHEM GQALCSVTQS LEKLQMLNAK**
651 KWREP

**b**

**Pkd1$^{KO}$:**

**Protein sequence coverage: 61%**

1 M**SELSDEASE PELLKR**SLSM WHGLGAQGSP EELDVPLDLH TAASIGQHEV
51 VKECVQR**GEL DLNKK**NGGGW TALMYASYIG HDTIVHLLLE AGVSVNVPTP
101 EGQTPLMLAS SCGNESIAYF LLQQGAELEM KDIHGWTALF HCTSAGHQQM
151 VK**FLLESGAN ANVR**EPVYGY TPLMEAAASG HEIIVQYFLN HGVKVDTR**DH**
201 **SGATACMLAR** QFGHMK**IVAL METHSPVLPK** SLYR**SPEKYE DLSSSDESWP**
251 **VPQR**QRPCRK **KGLSIHEGPR** ALAK**ITAIGL GGKTQTTYEQ VPPRGYVTFT**
301 **SSDENTMESE GLCYRDVTSP INEQDVESSS SSSREEPTFC ASLGPVWRSS**
351 **SSDGLARAQG LSSEASIESN EDSDHARK**SS VRKQTRSYLK NKSR**HGNSDG**
401 **HWPSSTGPAS IPGSEPQTEK SPYSGPQDLA TLLEQIGCLK YLQVFEEQDI**
451 **DLRIFLTLTE SDLKEIGITL FGPK**RKMTSA IAR**WHSSARP PSDALELAYA**
501 **DRLETEMQEL AIQLHK**CCEE AEALR**GQVSQ EQELRAVVES CLLEQDSAR**K
551 DIHAQLQEAQ TLAQDAALVL DQLR**ACQAEL SAR**LRQHHSP R**EGTPNPHFL**
601 **SADSKGWPIP LQALSLPELS GALEDRVHEM GQALCSVTQS LEKLQMLNAK**
651 KWREP

**c**

**Anks3 pSer5:**

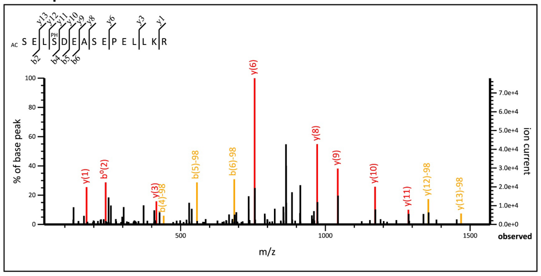


**Anks3 pSer2 & pSer5:**

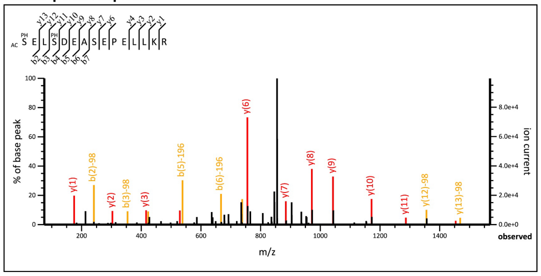


**Supplementary Figure 5. Identification of Anks3 phosphorylation sites. a,b**, LC-MS/MS peptide fragment coverage of Anks3 protein in WT and Pkd1$^{KO}$ cells. Matched peptides were shown in bold red. **c**, LC-MS/MS spectra for Ser5 single-phosphorylated Anks3 peptide and Ser2/Ser5 dual-phosphorylated Anks3 peptide.

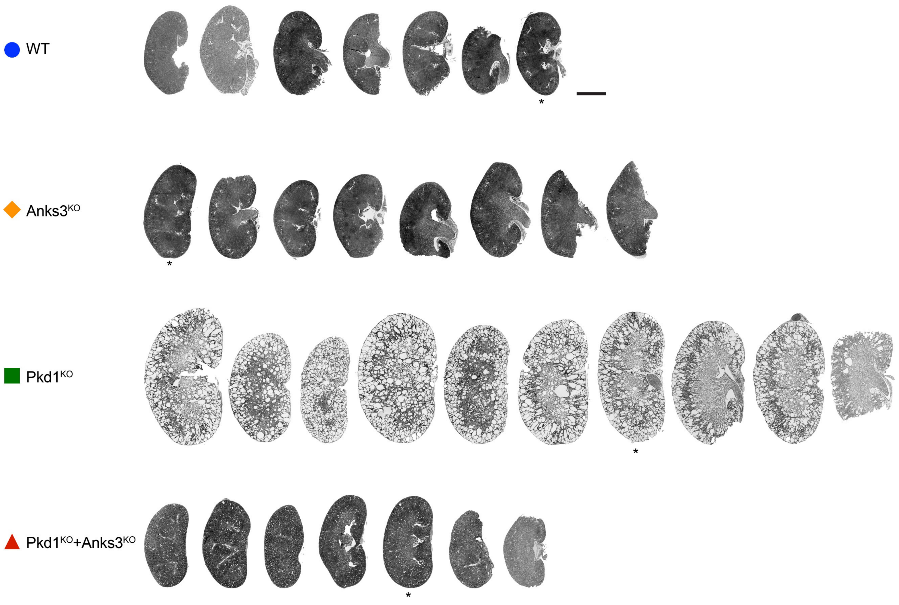


**Supplementary Figure 6. Images of all the histological sections used in Figure 3a-e.** Scale bar, 2 mm. Asterisks indicate the representative images used in Figure 3a.

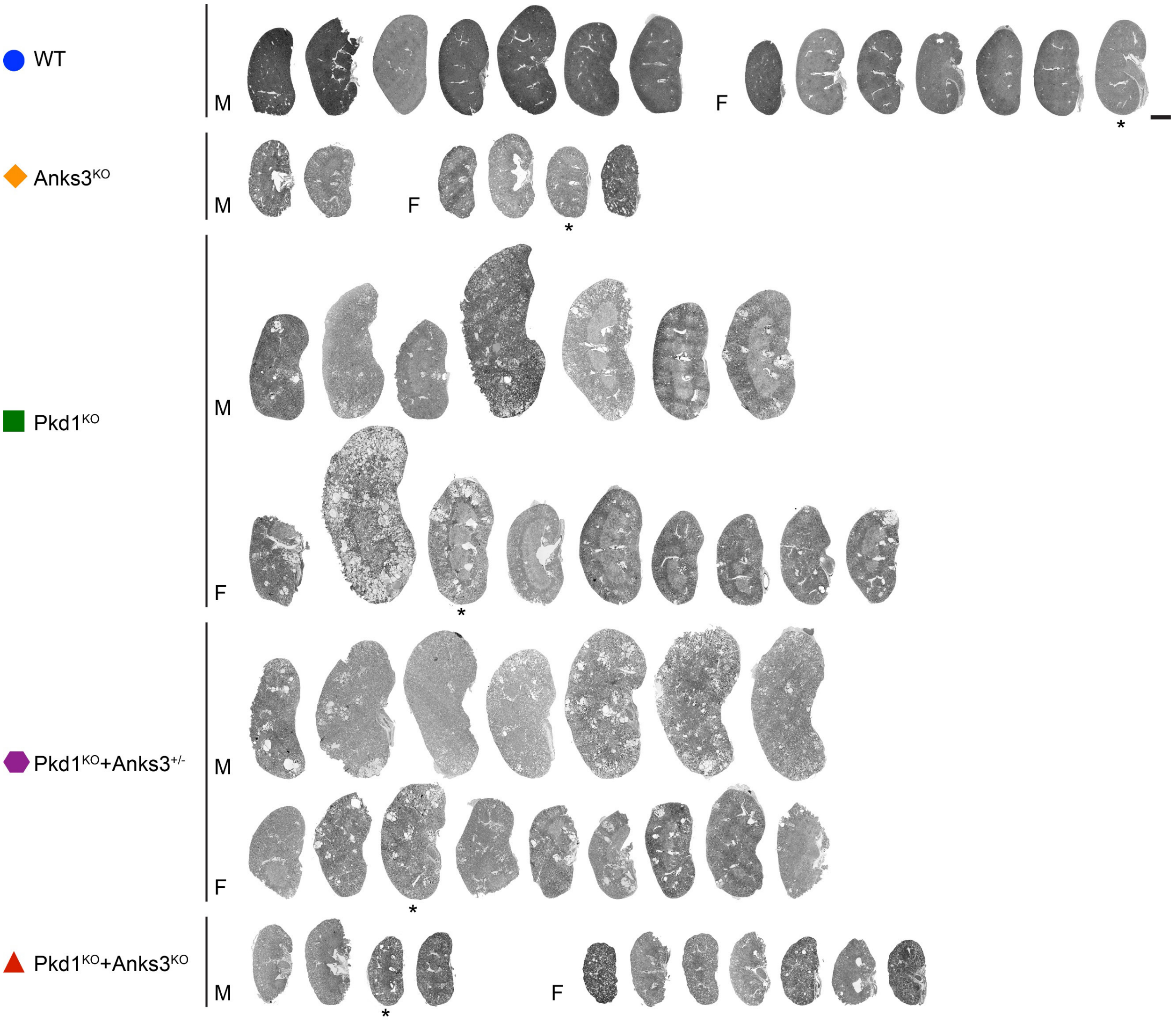


**Supplementary Figure 7. Images of all the histological sections used in Figure 3h-l.** Scale bar, 2 mm. Asterisks indicate the representative images used in Figure 3h.

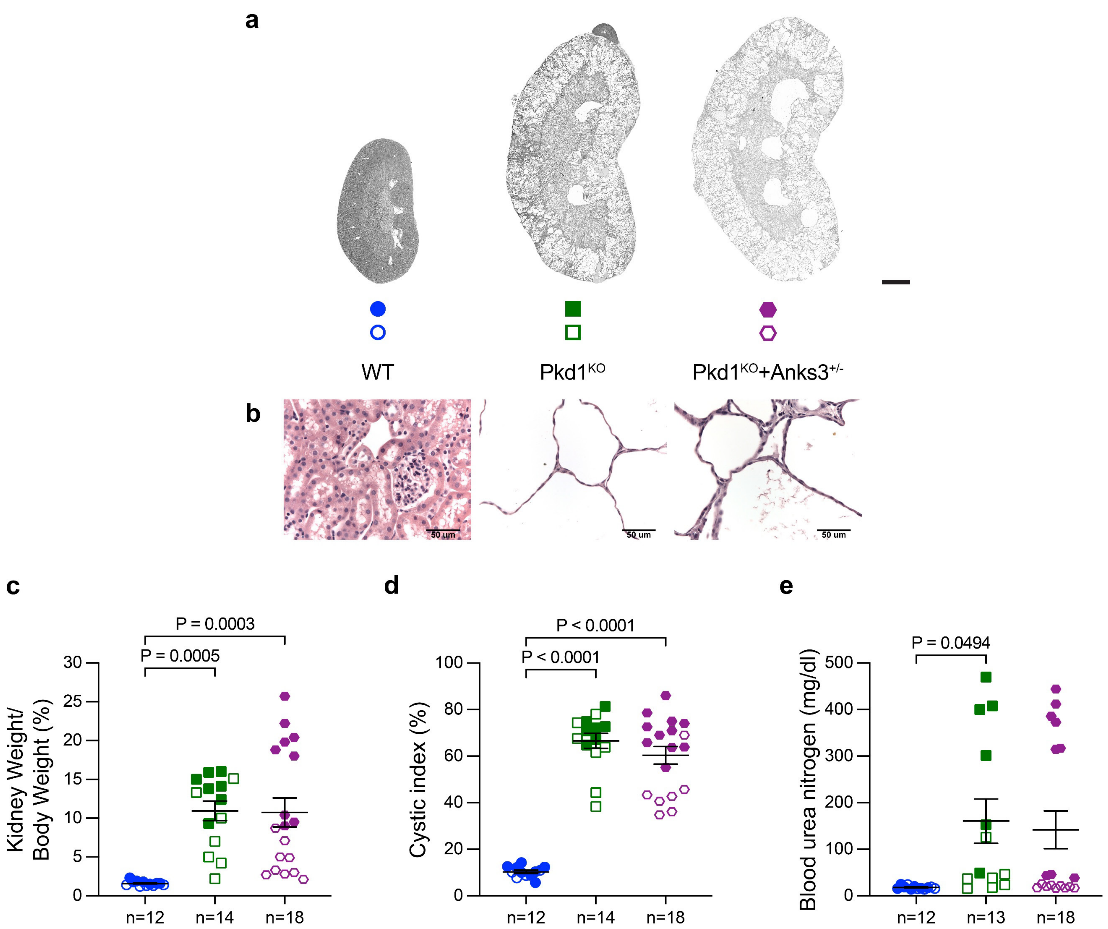


**Supplementary Figure 8. Heterozygous inactivation of Anks3 is not sufficient to suppress polycystic kidney disease in adult models.** **a**, Representative images of kidneys from mice at 19 weeks of age with indicated genotypes. All mice were administrated oral doxycycline from P28-P42 and were examined at 19 weeks. Scale bar, 2 mm. **b**, Representative images of H&E staining for the corresponding genotype. Scale bar, 50 μm. **c-e**, Aggregate quantitative data for kidney-to-body weight ratio (**c**), cystic index (**d**), and blood urea nitrogen (**e**). Colors and symbol shapes correspond to genotype defined in **a**; closed symbols, male; open symbols, female. *n*, number of mice in each group. Multiple-group comparisons were performed by one-way ANOVA followed by Tukey's multiple-comparison test, presented as mean ± s.e.m.

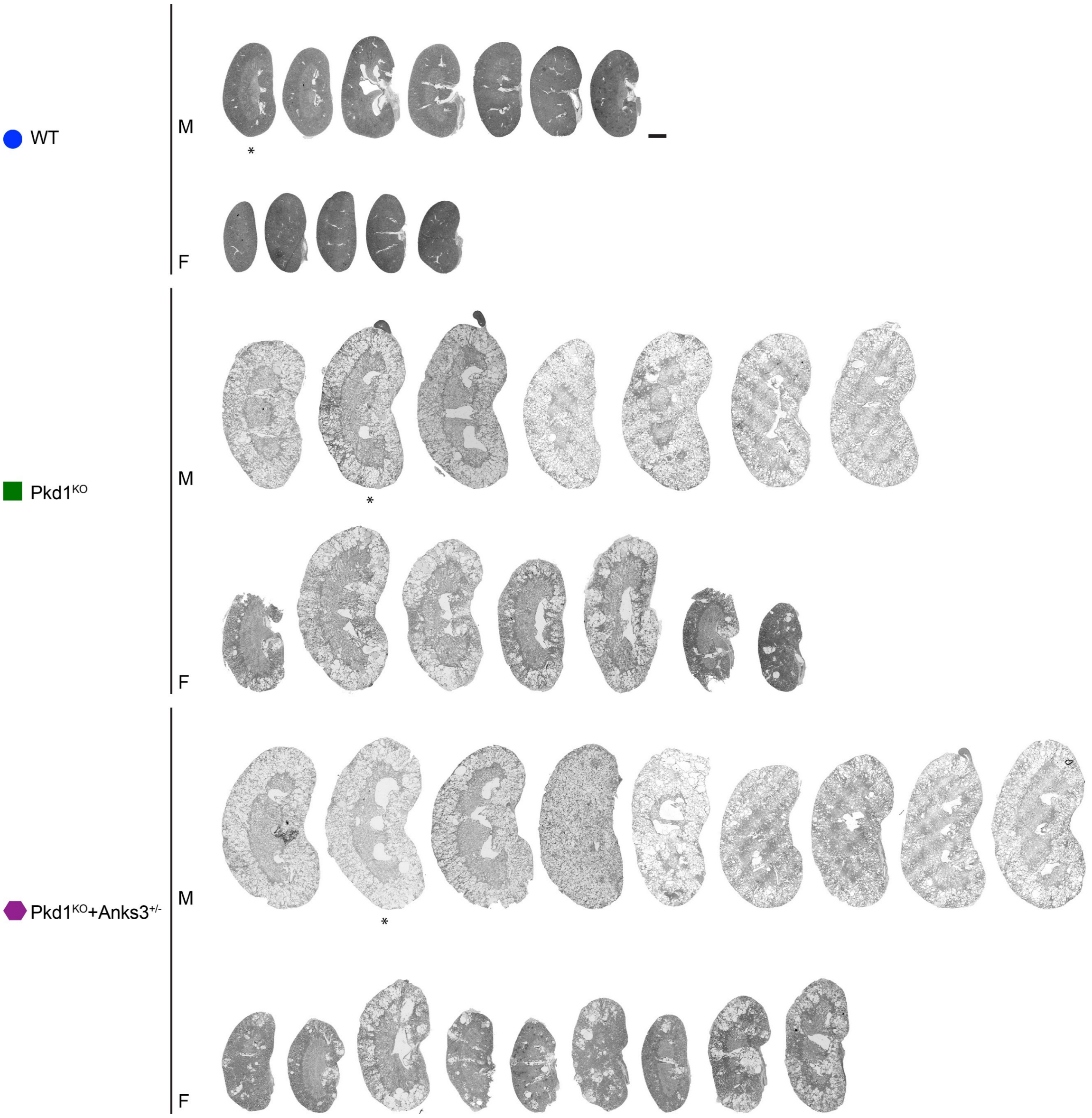


**Supplementary Figure 9. Images of all the histological sections used in Supplementary Figure 8.** Scale bar, 2 mm. Asterisks indicate the representative images used in Supplementary Figure 8a.

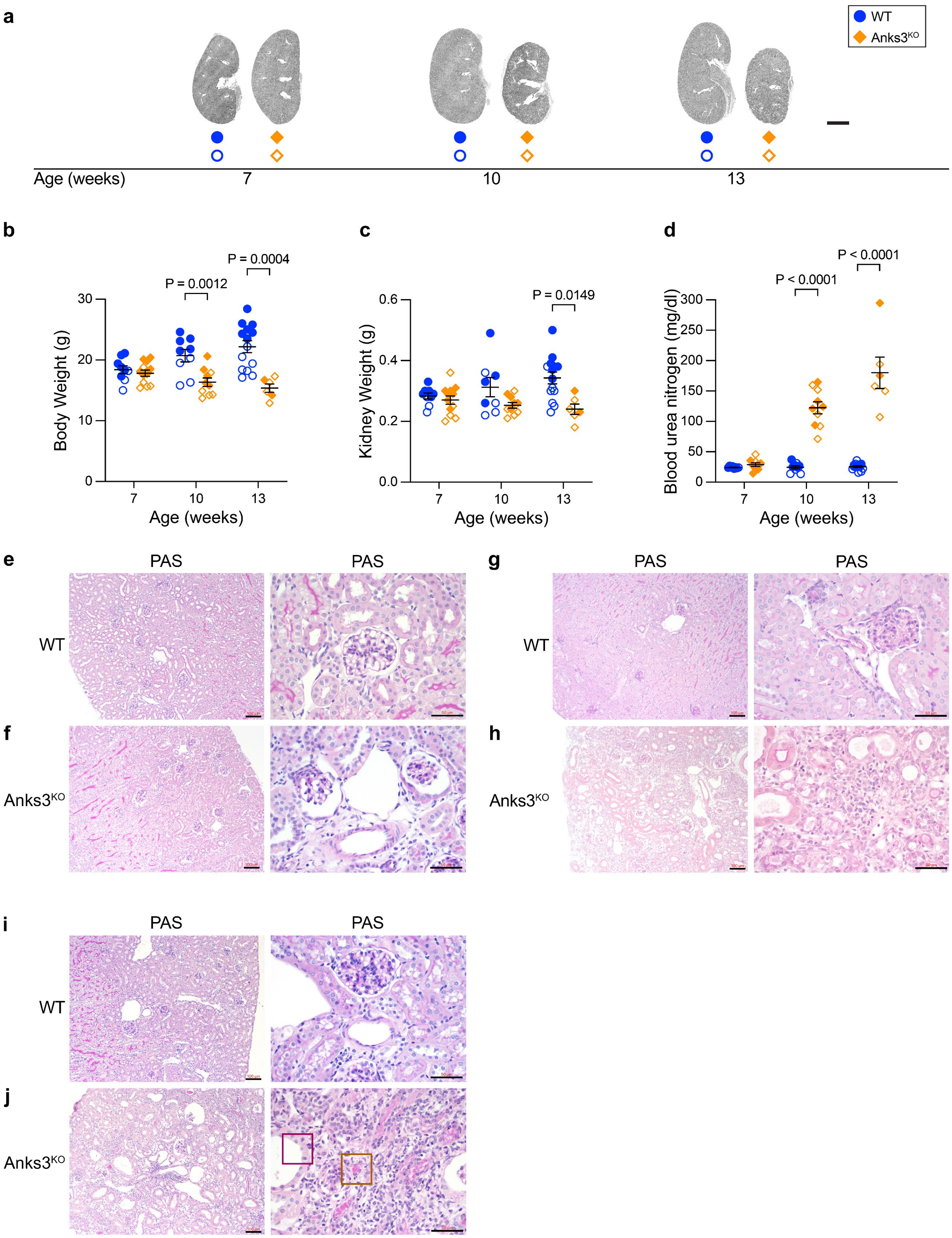


**Supplementary Figure 10. Kidney selective inactivation of Anks3 results in progressive kidney damage. a**, Representative images of kidneys from mice with the indicated ages and genotypes. All mice were administrated with oral doxycycline from P28-P42, and examined at either 7, 10, or 13 weeks to show the progression of the phenotype. The representative images of 13 weeks are reused from Figure 3h for the purpose of comparison. Scale bar, 2 mm. **b-d**, Aggregate quantitative data for body weight (**b**), kidney weight (**c**), and blood urea nitrogen (**d**). Colors and symbol shapes correspond to genotype defined in **a**, with indicated ages; closed symbols, male; open symbols, female. Blood urea nitrogen data at 13 weeks are reused from Figure 3l for the comparison purpose. Statistical significance by unpaired two-tailed Student's *t* test presented as mean ± s.e.m. **e-j**, Representative images of periodic acid-Schiff (PAS) stained kidney histological sections (*left*, low and *right*, high magnification) with the indicated genotypes. **e,f**, 7-weeks; **g,h**, 10-weeks; **i,j**, 13-weeks. Scale bar, 100 μm and 50 μm.

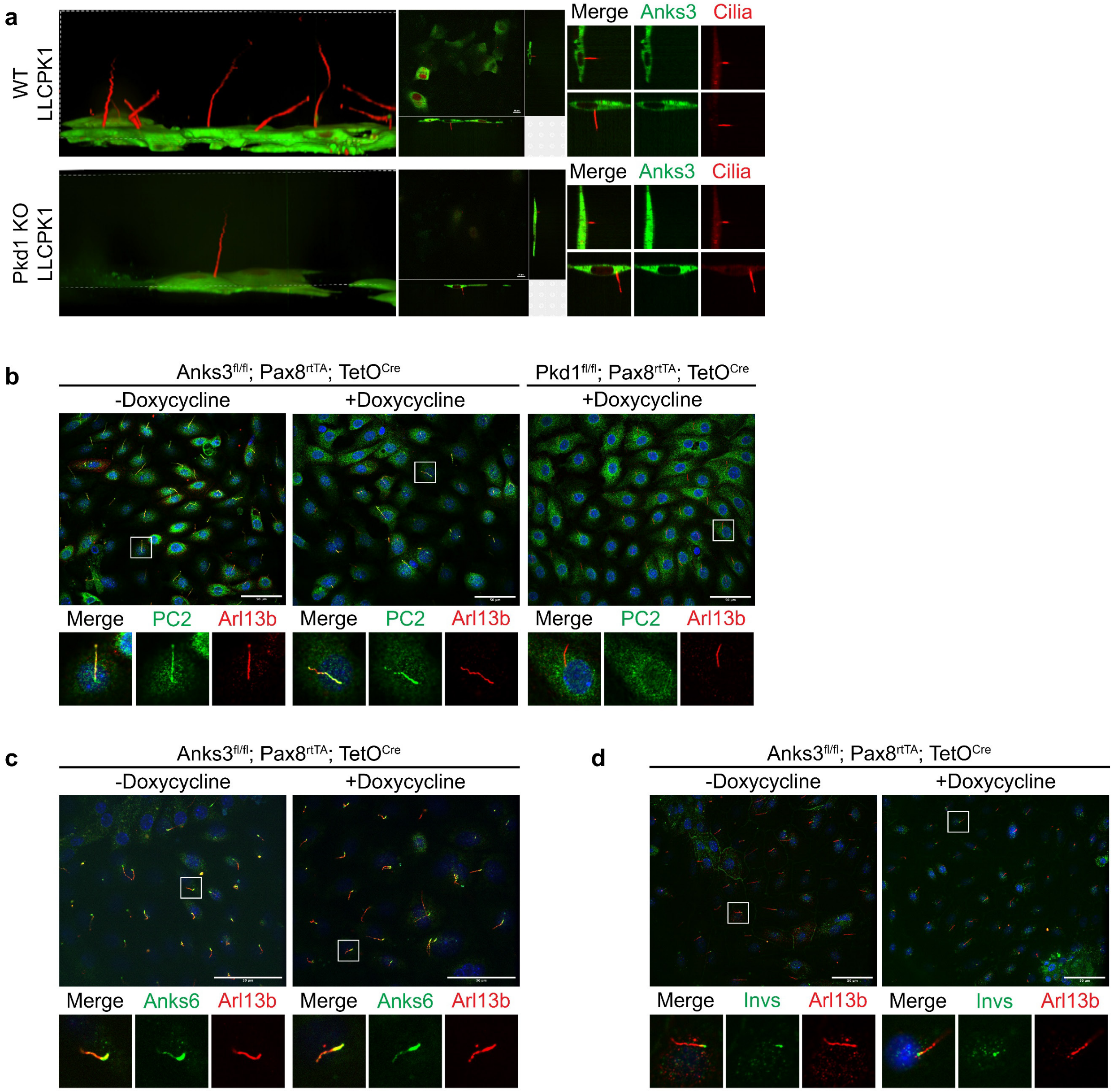


**Supplementary Figure 11. Inactivation of Anks3 does not affect localization of ciliary proteins. a**, Anks3 is not expressed in cilia. 3D reconstruction and z-sections for live cell confocal imaging of WT or Pkd1$^{KO}$ LLC-PK$_1$ cells stably expressing bicistronic Anks3-EGFP-PT2A-Nphp3$^{1-200}$-mApple. Scale bar, 10 μm. **b-d**, Anks3 inactivation does not change cilia localization of Arl13b, PC2 or its interacting proteins Anks6 and Invs. Double indirect immunofluorescence of PC2 (**b**), Anks6 (**c**) or Invs (**d**) and the cilia marker Arl13b (**b-d**) on ciliated primary kidney cell cultures from mice with the indicated genotypes, with or without in vitro doxycycline treatment. Scale bar, 50 μm.

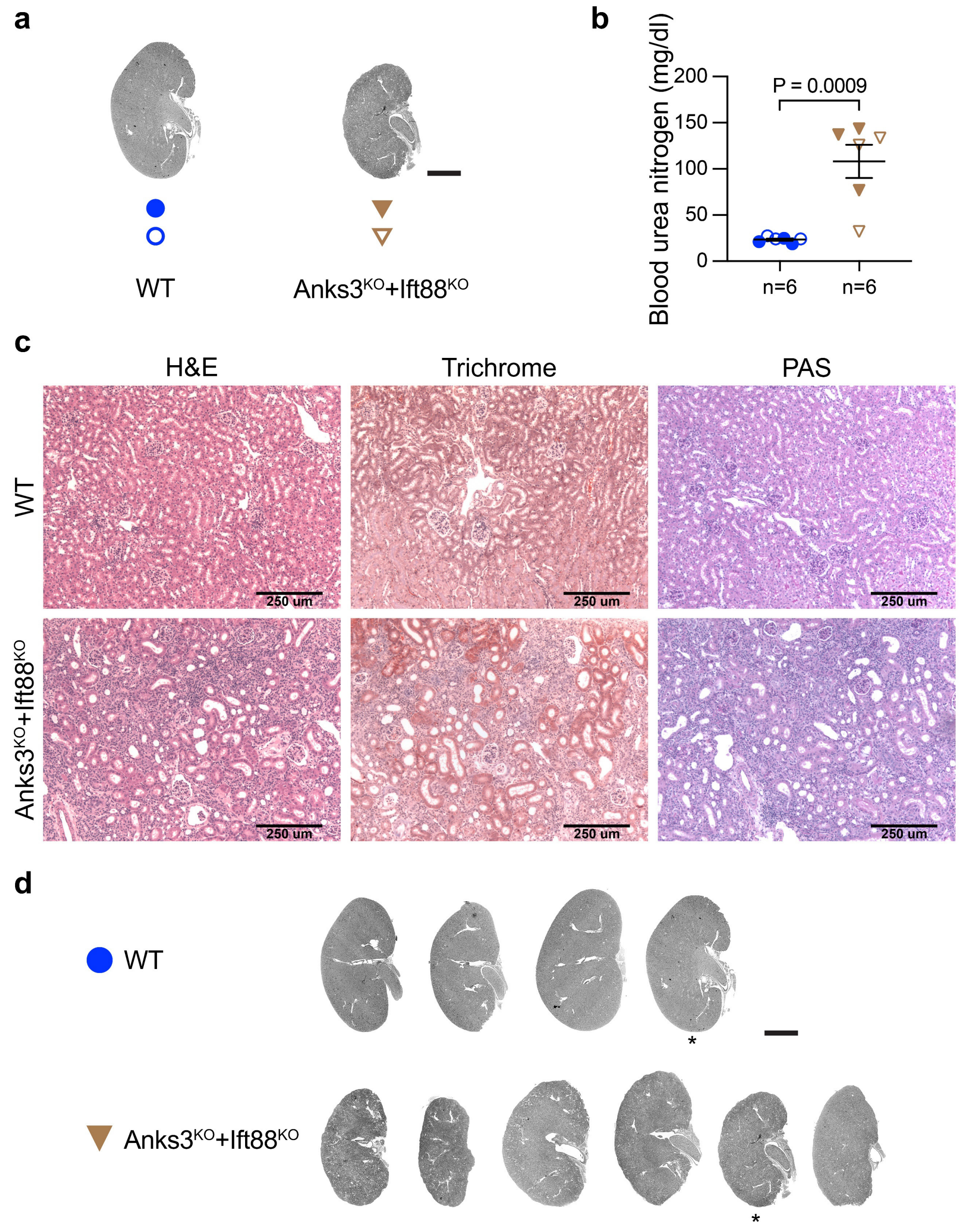


**Supplementary Figure 12. The kidney phenotype of Anks3 inactivation is not cilia dependent. a**, Representative images of kidneys from mice at 10 weeks of age with indicated genotypes. All mice were administrated with oral doxycycline from P28-P42. Scale bar, 2 mm. **b**, Aggregate quantitative data for blood urea nitrogen. Colors and symbol shapes correspond to genotype defined in **a**; male, closed symbols; female, open symbols. *n*, number of mice in each group. Statistical significance was determined by unpaired two-tailed Student's *t* test and presented as mean ± s.e.m. **c**, Representative images of H&E, Masson-t richrome, and PAS staining for the corresponding genotype. Scale bar, 250 μm. **d**, Images of all the histological sections used in **a**. Scale bar, 2 mm. Asterisks indicate the representative images used in **a**.

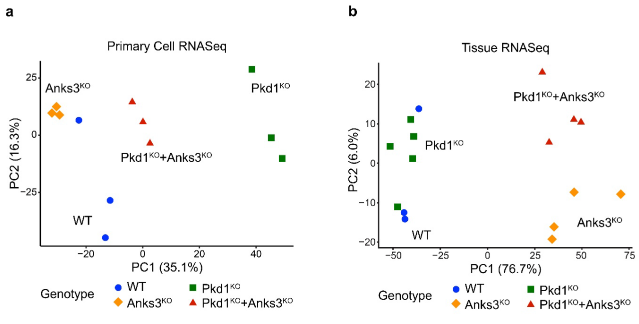


c

| Primary Cell RNASeq | FDR ≤ 0.05 | FDR ≤ 0.01 | |
|---|---|---|---|
| Anks3$^{KO}$ *vs* WT | 1347 | 911 | 362 / 549 |
| Pkd1$^{KO}$ *vs* WT | 3831 | 2833 | 1571 / 1262 |
| Pkd1$^{KO}$+Anks3$^{KO}$ *vs* WT | 2009 | 1406 | 640 / 766 |
| Pkd1$^{KO}$+Anks3$^{KO}$ *vs* Anks3$^{KO}$ | 3366 | 2376 | 1276 / 1100 |
| Pkd1$^{KO}$+Anks3$^{KO}$ *vs* Pkd1$^{KO}$ | 3311 | 2319 | 943 / 1376 |

d

| Tissue RNASeq | FDR ≤ 0.01 | FDR ≤ 0.01 $\lvert log_2FoldChange\rvert>1$ | |
|---|---|---|---|
| Anks3$^{KO}$ *vs* WT | 8205 | 2482 | 1790 / 692 |
| Pkd1$^{KO}$ *vs* WT | 737 | 110 | 65 / 45 |
| Pkd1$^{KO}$+Anks3$^{KO}$ *vs* WT | 7414 | 1889 | 1436 / 453 |
| Pkd1$^{KO}$+Anks3$^{KO}$ *vs* Anks3$^{KO}$ | 1476 | 185 | 112 / 73 |
| Pkd1$^{KO}$+Anks3$^{KO}$ *vs* Pkd1$^{KO}$ | 8069 | 2193 | 1623 / 570 |

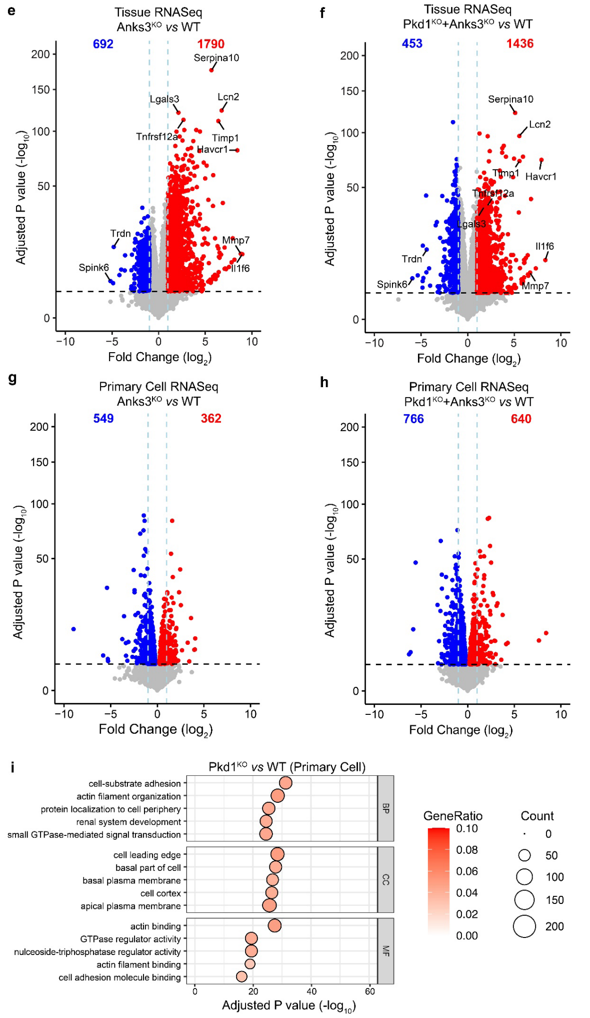


**Supplementary Figure 13. RNASeq from allelic series of primary kidney cell cultures and bulk kidney tissues. a,b**, PCA plots of RNASeq from primary cells (**a**) and bulk kidney tissues (**b**) showing the global transcriptomic difference among the indicated genotypes. **c,d**, Tables showing the number of differentially expressed genes (DEGs) in primary cell (**c**) and tissue (**d**) RNASeq with the indicated pairwise comparison and threshold criteria. Last column shows a breakdown for the number of upregulated (red) and downregulated DEGs (blue). **e-h**, Volcano plots of DEGs in tissue (**e,f**) and primary cell (**g,h**) RNAseq in the two pairwise comparisons: Anks3$^{KO}$ *vs* WT (**e,g**) and Pkd1$^{KO}$+Anks3$^{KO}$ *vs* WT (**f,h**). Genes with significant differential expression are indicated by red (upregulated) and blue (downregulated) dots. Black dashed line, FDR $\leq$0.01 threshold; blue dashed line, two-fold change threshold. The most significantly altered genes and the genes with largest absolute fold change in **e** are labeled in **f**. **i**, Gene ontology analysis performed using the *clusteProfiler* R package for Pkd1$^{KO}$ *vs* WT in primary cells with FDR $\leq$0.01 threshold.
